# BAX and SMAC Regulate Bistable Properties of the Apoptotic Caspase System

**DOI:** 10.1101/867663

**Authors:** Stephanie McKenna, Lucía García-Gutiérrez, David Matallanas, Dirk Fey

## Abstract

The initiation of apoptosis is a core mechanism in cellular biology by which organisms control the removal of damaged or unnecessary cells. The irreversible activation of caspases is essential for apoptosis, and mathematical models have demonstrated that the process is tightly regulated by positive feedback and a bistable switch. BAX and SMAC are often dysregulated in diseases such as cancer or neurodegeneration and are two key regulators that interact with the caspase system generating the apoptotic switch. Here we present a mathematical model of how BAX and SMAC control the apoptotic switch. Formulated as a system of ordinary differential equations, the model summarises experimental and computational evidence form the literature and incorporates the biochemical mechanisms of how BAX and SMAC interact with the components of the caspase system. Using simulations and bifurcation analysis, we find that both BAX and SMAC regulate the time-delay and activation threshold of the apoptotic switch. Interestingly, the model predicted that BAX (not SMAC) controls the amplitude of the apoptotic switch. Cell culture experiments using siRNA mediated BAX and SMAC knockdowns this model prediction. We further validated the model on data of the NCI-60 cell line panel using BAX protein expression as cell-line specific parameter and show that model simulations correlated with the cellular response to DNA damaging drugs and established a defined threshold for caspase activation that could distinguish between sensitive and resistant melanoma cells. In summary, we present an experimentally validated dynamic model that summarises our current knowledge of how BAX and SMAC regulate the bistable properties of irreversible caspase activation during apoptosis.

## Introduction

Correct regulation of cellular homeostasis is fundamental in multicellular organisms. Thus, this process is tightly regulated by complex molecular networks which control cell fate, determining when a cell should proliferate, differentiate or die through programmed cell death. Apoptosis is an essential form of programmed cell death which mediates the removal of damaged or unnecessary cells. Tight regulation of apoptosis is vital to the maintenance of normal cell function. Apoptosis can proceed via the extrinsic pathway, induced by binding of an extracellular ligand to its death receptor, or the intrinsic pathway, activated by a diverse array of cytotoxic stimuli which converge on the outer mitochondrial membrane^1,2^. Subsequent release of pro-apoptotic proteins from the mitochondria is regulated by pro-apoptotic and anti-apoptotic BCL2 proteins^2^. Both apoptotic pathways are mediated by the cleaving proteins, caspases, which are produced as inactive zymogens^3^. Initiator caspases (2, 8, 10, 9) are activated by pro-apoptotic proteins and subsequently cleave and activate executioner caspases (3, 6, 7) which act directly on cell dismantling substrates^2,3^. Apoptosis is triggered when the caspase cascade is irreversibly activated, and several proteins regulate the correct switching on of this machinery to ensure avoidance of incomplete cell death. Positive and negative regulators of the caspases include the BCL2 family of proteins^4–7^, the inhibitor of apoptosis (IAP)^8–10^ family and several mitochondrial proteins such as APAF^11^, Cytochrome C (CytC)^5,7,12^ and SMAC (Second mitochondria-derived activator of caspases), also called Direct IAP-binding Protein with low pI (DIABLO^5,12–15^, hereto SMAC). Importantly, dysregulation of apoptosis is common in disease, and is a hallmark of cancer and neurodegenerative diseases^16,17^.

Given its central role in life, the mechanisms that regulate apoptotic networks are the focus of intensive research. This work has shown that there are several control points which determine the activation of apoptosis upon pro-apoptotic signals^5^. Given the complex dynamics regulating these mechanisms, several groups have applied systems biology approaches to study the mechanistic regulation of these apoptotic networks^18–24^. It has been shown that the relative stoichiometry of the different proteins, caspase cleavage and protein degradation rates determine the commitment of the apoptotic machinery to cell death. In the case of the intrinsic pathway it has been experimentally and computationally demonstrated that caspase dependent apoptosis is a highly controlled process which exhibits an irreversible, bistable switch^18–20,25^. Mathematically, bistability is the coexistence of two stable steady states, allowing for rapid, often irreversible transition between these steady states. Once a stimulus threshold is exceeded, the system can switch from a low level of activation (the ‘off’ stable steady state) to a high level of activation (the ‘on’ stable steady state) in an all-or-nothing fashion^20^. Phenomenologically, a bistable switch exhibits two characteristic properties. On the level of the dose-response: the emergence of the hysteresis effect which implies that for the system to switch “on” from one steady state to the other, the input stimulus must exceed a certain threshold. The system will reside in this new steady state, even if the stimulus changes. In order to switch “off” to the preceding steady state, a different, lower stimulus threshold must be reached. The hysteresis effect indicates that the system maintains different on and off thresholds. On the level of the time-course, a characteristic time-delay precedes the rapid switch-like transition to the other stable steady-state. Upon application of the input stimulus, the system remains relatively unchanged in the “off” state for some amount of time before rapidly activating in a snap-like fashion. This time-delay of activation can be increased or decreased by alteration of the stimulus concentration or modification of model components.

The primary mechanism generating bistability in the intrinsic pathway is positive feedback^20^. This positive feedback is realised by an explicit and implicit feedback loop. Explicitly, upon cleavage and activation of caspase 3 by caspase 9, caspase 3 can in turn cleave and activate caspase 9^20^. In the implicitly hidden feedback loop, IAPs can bind and inhibit all caspases within the cascade. Caspase 3 relieves inhibition of its upstream activator caspase 9 through IAP binding, thereby creating a second, implicit positive feedback loop to contribute to the bistability of the system^20^. Previous work indicates that the explicit and implicit feedback mechanisms could be an important feature in determining how the apoptotic system achieves ultra-sensitivity, bistability and irreversibility. In order to get a better understanding of the properties of the bistable switch in the intrinsic apoptotic pathway and understand how its deregulation may be related to disease it is necessary to develop models that include other regulatory proteins of the network. Models developed including BAX and SMAC have shown that these proteins are key in the regulation of apoptotic commitment. BAX is the gateway to mitochondrial activation of intrinsic apoptosis and has been shown to form pores in the mitochondrial membrane resulting in the release of pro-apoptotic proteins^26–28^. SMAC is one of the proteins released from the mitochondria upon activation of the intrinsic apoptotic pathway and is and inhibitor of IAPs^13^. Correct function of both BAX and SMAC is central to the regulation of apoptosis. Importantly, deregulation of these pro-apoptotic proteins has been described in tumours and patients with neurodegenerative diseases^4,6,26,28–42^.

Here, we wanted to understand how BAX and SMAC control the emergent properties of the bistable switch. Thus, we have built a mathematical model of BAX and SMAC interactions with the caspase system which mirrors experimental observations from our own experiments and the literature. Here we show that once the caspase 3 activation threshold is reached, caspase 3 switches to the high activity ‘on’ state in an irreversible, all-or-nothing manner, which is in line with previously published models^18–20,22,43^. We demonstrate that upon increasing the input, BAX expression level or SMAC release rate were shown to reduce the time lag in caspase 3 activation. Our bifurcation analyses revealed that an increase in BAX expression resulted in a reduction in the caspase 3 activation threshold. An increase in SMAC release rate also exhibited a reduction in caspase 3 activation threshold, though to a lesser extent. Simulations confirmed by validation experiments show that the caspase system might be differentially sensitive to BAX and SMAC mimetics. Finally, we further validated the model on data of the NCI-60 cell line panel using BAX protein expression as cell-line specific parameter. The model simulations correlated with the cellular response to DNA damaging drugs, indicating that our model can be used for in silico simulations of pathological changes of the apoptotic machinery.

## Results

### Model construction

#### Dynamic modelling of BAX and SMAC regulated intrinsic apoptotic pathway

To elucidate how BAX and SMAC regulate the bistable switch within the intrinsic apoptosis pathway, we constructed a mathematical model of Ordinary Differential Equations (ODEs). To construct the model, we extended the model of caspase activation by Legewie et al.^20^ with information from the literature to include (i) BAX mediated initiation of the caspase cascade and (ii) SMAC-mediated sequestration of IAP^44–46^. The reaction kinetic scheme of the model is outlined in Figure 1. The expansion of the model to include BAX and SMAC is detailed below.

**Figure 1.**
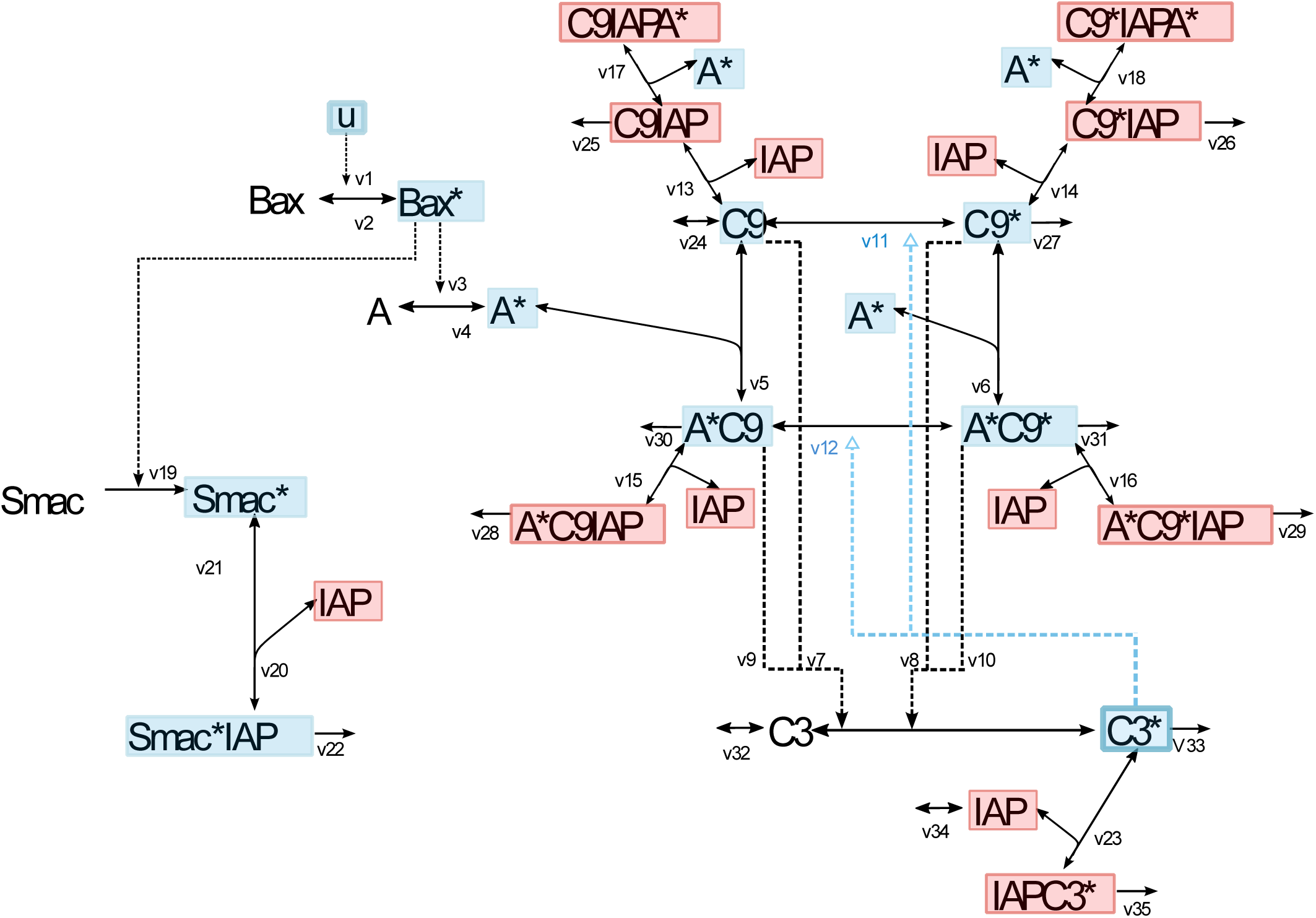
Reaction kinetic scheme of the intrinsic apoptosis model. A pro-apoptotic insult, u, activates BAX. Upon activation of BAX, heptamerisation of Apaf-1 (A) occurs to form the active apoptosome (A*), which by recruitment of caspase 9 forms complexes that can activate the intrinsic apoptosis cascade. A* can recruit and stimulate both pro-caspase 9 (C9) and cleaved caspase 9 (C9*). When active, caspase 9 causes cleavage and activation of pro-caspase 3 (C3). In turn, active caspase 3 (C3*) can activate caspase 9 through a positive feedback loop. Further, inhibitor of apoptosis protein (IAP) can bind to all forms of caspase 9 and caspase 3 and block their activity. Active SMAC* is released by Bax* and can bind the free form of IAP leading to the degradation of the SMAC*IAP complex. * denotes the active form of the protein

In the Legewie model, caspase activation is initiated by apoptosome (denoted A* in the model, Figure 1) formation through heptamerisation of APAF-1 and binding to procaspase 9 resulting in procaspase 9 autoproteolysis and activation. Active procaspase 9 (C9) then causes caspase 9 (C9*) cleavage and activation. Data has shown that auto-proteolysis of C9 does not impact its recruitment to A*^47,48^, and so, it has been assumed that A* can recruit and stimulate both C9 and C9*. C9* will activate procaspase 3 (C3) which can in turn cleave and activate caspase 3 (C3*) to act directly on cell dismantling substrates resulting in cell death. C3* can activate caspase 9 through a positive feedback loop. IAP (X), which maintains an E3 ligase domain, can reversibly bind and inhibit all forms of caspase 9 and caspase 3 ^15^ to prevent the activation of cell death.

A limitation of this model is that it does not consider evidence that shows that Cytochrome C (CytC) release from the mitochondria is dependent on BAX pore formation^7^ and further that CytC release itself is insufficient to completely prevent the IAP’s inhibitory effect upon caspases which requires the participation of SMAC^49^. Thus, the model was expanded to include apoptotic regulators, BAX and SMAC. BAX becomes activated (BAX*) by a pro-apoptotic insult (u). To keep the model simple the depolarisation of the mitochondrial membrane by BAX, the release of CytC from the mitochondria and the subsequent activation of APAF-1 causing formation of the apoptosome (A*), was lumped into a single reaction in which A* formation is dependent on active BAX. Concomitantly, SMAC is released from the mitochondria in a BAX* dependant manner to bind and form an inhibitory complex with IAP, which can be degraded^14,50^. More details about the modelling of BAX and SMAC are provided in two dedicated sections below.

A full account of all reactions, rate laws, parameters and equations, including initial conditions is given in Tables 1 and 2. This ODE model provides a mathematical representation to describe exactly how a pro-apoptotic insult activates the intrinsic apoptosis pathway and how these species change over time. The model can be used to simulate how cell-line and patient-specific differences of gene and protein expression may alter the system dynamics of caspase activation - predictions which may be experimentally validated.

**Table 1.**
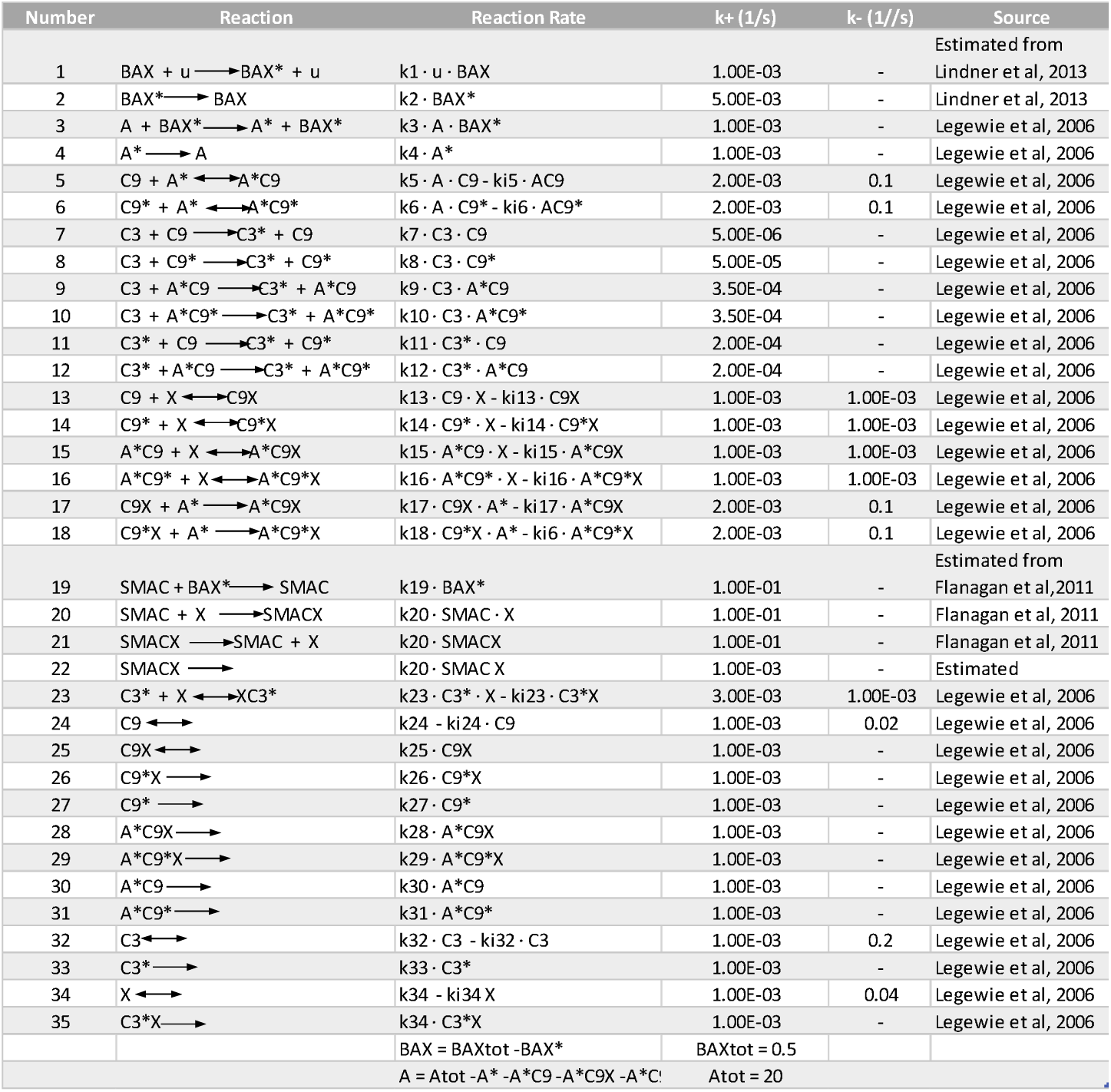
Table of Model Reactions and Reaction Rates.

**Table 2.** Table of Ordinary Differential Equations.

| Left-hand Sides | Right-hand Sides | Initial Concentrations (nM) |
| --- | --- | --- |
| $d[BAX^*]/dt$ | $v1 - v2$ | 0 |
| $d[A^*]/dt$ | $v3 - v4 - v5 - v6 - v17 - v18$<br>$+v28 + v30 + v31 + v29$ | 0 |
| $d[C9]/dt$ | $-v5 - v11 - v13 + v24$ | 20 |
| $d[C9X]/dt$ | $v13 - v17 - v25$ | 0 |
| $d[X]/dt$ | $-v13 - v14 - v15 - v16 - v20$<br>$+v21 - v23 + v34$ | 20 |
| $d[A^*C9X]/dt$ | $v15 + v17 - v28$ | 0 |
| $d[A^*C9]/dt$ | $v5 - v15 - v30$ | 0 |
| $d[C3]/dt$ | $-v7 - v8 - v9 - v10 + v32$ | 200 |
| $d[C3^*]/dt$ | $v7 + v8 + v9 + v10 - v23 - v33$ | 0 |
| $d[C3^*X]/dt$ | $v23 - v35$ | 0 |
| $d[C9^*X]/dt$ | $v14 - v18 - v26$ | 0 |
| $d[C9^*]/dt$ | $v11 - v6 - v14 - v27$ | 0 |
| $d[A^*C9^*]/dt$ | $v6 + v12 - v16 - v31$ | 0 |
| $d[A^*C9^*X]/dt$ | $v16 + v18 - v29$ | 0 |
| $d[SMAC]/dt$ | $v19 - v20 + v21$ | 0 |
| $d[SMACX]/dt$ | $v20 - v21 - v22$ | 0 |

In the following section we review experimental evidence for the BAX and SMAC mechanisms of crosstalk with the intrinsic apoptosis pathway in detail and show how they are implemented in the dynamic model. Finally, we explore the intricate kinetic behaviour and dynamics of BAX and SMAC-regulated caspase activation.

#### Modelling BAX

As already mentioned, as one of the most important proteins in the BCL2 pro-apoptotic family, BAX mediates the activation of intrinsic apoptosis in response to DNA damage^51^.Upon pro-apoptotic insult, BAX is translocated to the mitochondrial membrane where it can homo-oligomerise to form the ‘apoptotic pore’ within the membrane^52^. This mitochondrial outer membrane permeabilization (MOMP) results in the release of CytC from the mitochondria, triggering activation of the caspase cascade^52^. Pro-apoptotic BCL2 proteins including BAX can be deactivated by binding of antiapoptotic BCL2 proteins, thereby blocking activation of the intrinsic apoptosis cascade^52^.

In our model BAX acts upstream of APAF-1 heptamerisation (Figure 1). BAX activation and translocation are modelled in a single reaction (reaction v1, Figure 1). Neglecting biological details such as the inhibitory binding and neutralisation of BCL-XL to BAX^7,37^, deactivation of BAX* is modelled as a simple first order reaction (v2, Figure 1). Active BAX* subsequently mediates the formation of the apoptosome, which is modelled as a first order reaction, dependant on BAX* (v3, Figure 1). A*subsequently triggers the activation of the caspase cascade.

#### Modelling SMAC

Following a pro-apoptotic insult and initiation of MOMP by the BAX mediated formation of pores in the mitochondrial membrane, SMAC is released from the mitochondria into the cytosol, where it binds and releases the inhibitory effect of IAP on caspases^13^. When SMAC binds the second or third BIR domain of IAP, its E3 ubiquitin ligase activity is stimulated, IAP is ubiquitinated and inhibited^53^. Degradation of IAP proteins following SMAC binding results in the loss of its inhibitory effect over procaspases 3 and 9^50^. In this way, SMAC induces the activation of procaspase 3 and mature caspase 3, thus promoting the induction of apoptosis^13^. Early evidence has shown that SMAC can exist as a dimer or monomer and that the dimer is essential for its pro-apoptotic role^13^. It was also shown that the N-terminal region of SMAC is necessary for the activation of caspase 3 and that a short peptide mimicking the N-terminus could mediate activation of apoptosis^13,49^, demonstrating the key role of this protein in the activation of the intrinsic apoptotic pathway. This has led to the development of a series of SMAC mimetics for the treatment of cancer that are being tested in the clinic^54^.

Based on this knowledge, and to include SMAC, we extended the core model by several processes. BAX* translocation to the mitochondria triggers MOMP-mediated SMAC release and dimerisation. To simplify the model, SMAC release and dimerisation was modelled as a single, first order reaction dependant on BAX* (v19, Figure1). Secondly, SMAC forms a complex with IAP to eliminate IAP inhibition of caspase activity (v20), relieving formation of inhibitory complexes with caspase 9 and caspase 3 (C9IAP, C9*IAP, A*C9IAP, A*C9*IAP, C3*IAP). We have considered SMACIAP complex formation to be a reversible reaction (v21). Degradation of the SMACIAP complex is represented by v22. We have made two assumptions in the modelling of SMAC dynamics. Firstly, we have assumed that the parameter values for SMAC release are similar to those of BAX dependant activation of A* as both are dependent on BAX mediated MOMP. Although, we have assumed SMAC release is 10 times faster than BAX activation of A* formation, to capture the extra time required for the multi-step mechanism of A* formation that was simplified in the model. Secondly, we have assumed that the parameter values for SMAC association with IAP are similar to the caspase-IAP interactions considering they bind at the same site^20^. Though, to ensure that SMAC has the ability to sequester IAP away from both caspase 3 and caspase 9 as has been experimentally observed^46^, we have modelled SMAC-IAP binding with slightly higher affinity.

#### The model simulates the experimentally observed timing and irreversibility of the apoptotic switch

First, we wanted to investigate if simulations of the model would mimic known experimental observations. Experiments carried out in cytosolic extracts have shown that upon exogenous application of CytC, maximal caspase 3 (C3*) activation (cleavage of procaspase 3) can occur within 15-30 minutes in some cells, depending on stimulus strength^11^. Protein concentrations of BAX (0.5nM), APAF-1 (20nM), caspase 9 (20nM), caspase 3 (200nM) and IAP (40nM) previously measured in HeLa cells were assumed^20,45^. A pro-apoptotic insult was stimulated by u=0.5nM. Simulation results in Figure 2A show that the time course of C3* activation agrees well with experimental findings. C3* activation occurs following 1500s (∼25 minutes) demonstrating that our model simulations mimic experimental observations^11^.

**Figure 2.**
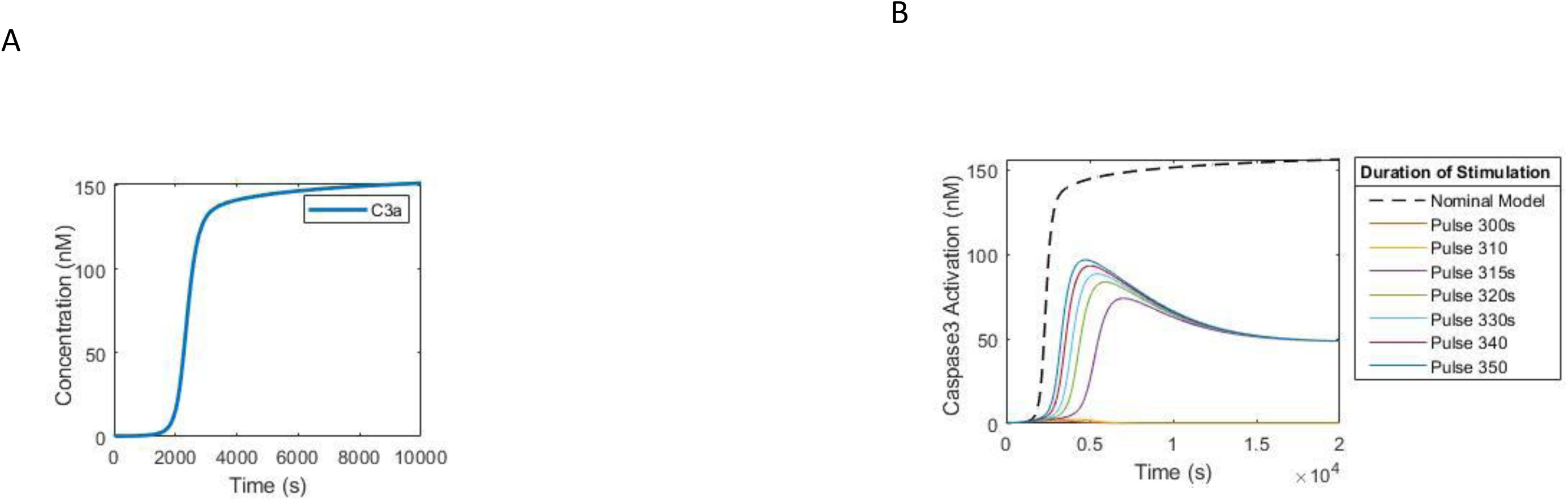
Investigating Model Behaviour. (A). Time course of activation of caspase 3 under constant stimulation. BAX was activated upon application of stimulus, *u*(0.5nM). The time course demonstrates that the model simulation of caspase 3 activation is in accordance with experimental results-that caspase 3 is activated within 15 minutes following initiation^14^. (B) Time course of caspase 3 activation in response to transient pulse stimulation. Upon stimulation of BAX by *u*(0.5nM) for 300 seconds, caspase 3 does not become activated. When a pulse stimulation of *u*(0.5nM) for 315 seconds was applied, caspase 3 was activated to ∼50% of nominal model caspase 3 activation. Once the stimulus is removed, caspase 3 remains activated demonstrating that once the stimulus threshold of 315 seconds is reached, caspase 3 activation is irreversible.

Apoptosis is generally considered to be irreversible once the executioner caspase 3 is activated beyond a certain threshold^55^, although it must be noted that recent evidence points to the reversibility of apoptosis in some cases^56^. Once caspase 3 is activated apoptosis can happen in a few minutes^57^. For this reason, we wanted to double check the irreversibility of the caspase-activation switch in our model^58^. We carried out a simulation study with differently sized pulse stimulations. Transient pulse stimulations with an amplitude of u=0.5nM and durations ranging 300s to 350s were used (Figure 2B). In the model, continuous stimulation of *u=*0.5nM activates C3* at 1500s. Transient pulse stimulation for 310s (*u=*0.5nM) does not result in activation of C3*. But when *u=*0.5nM is applied for 315s, C3* activation switches on to reach ∼50% of C3* the activation level for continuous stimulation. Increasing lengths of stimulus pulses (320-350s) increases the C3* activation level. Transient pulse activation of the system results in a lower C3* activation level when compared to the nominal model. However, it is important to note that once the C3* activation threshold of 315s (∼5mins) stimulation by *u=*0.5nM is reached, C3* activation is irreversible. These results demonstrate that the model that we have developed reflects the experimental observations that apoptosis is irreversible once the activation threshold of caspase 3 is reached. Moreover, they also demonstrate that as shown experimentally^57^, caspase activation is regulated by the duration (Figure 2B) and strength (Figure 4A) of the pro-apoptotic signal.

**Figure 3.**
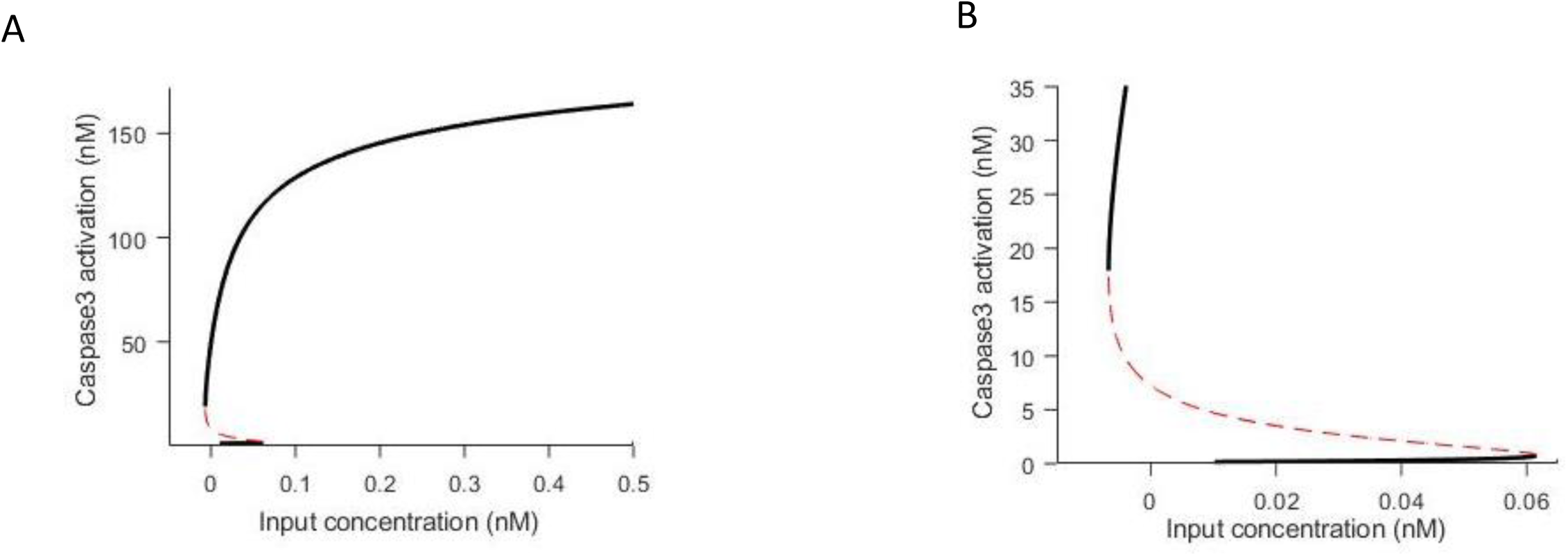
Bistable Behaviour of Caspase 3 activation. Steady state stimulus response bifurcation curve of caspase 3 activation. Figure 3 displays that simulated caspase 3(C3*) activation demonstrates irreversible bistability and hysteresis. (A). Three steady states are exhibited by the system, two stable (solid line) and one unstable (dashed line). The system can reside in one of the two stable steady states (‘on’ or ‘off’) but not in the intermediate unstable steady state. (B). The first bifurcation point at 0.06 and the second at −0.01 indicates that the system displays hysteresis between stimulus concentration (*u*) 0-0.06nM. The system remains at low caspase 3 activity (‘off’ state) for increasing stimuli until a threshold of *u* (0.06 nM) is reached, at which point caspase 3 activity switches to the second steady state (‘on’ state) irreversibly, in an all-or-none fashion.

**Figure 4.**
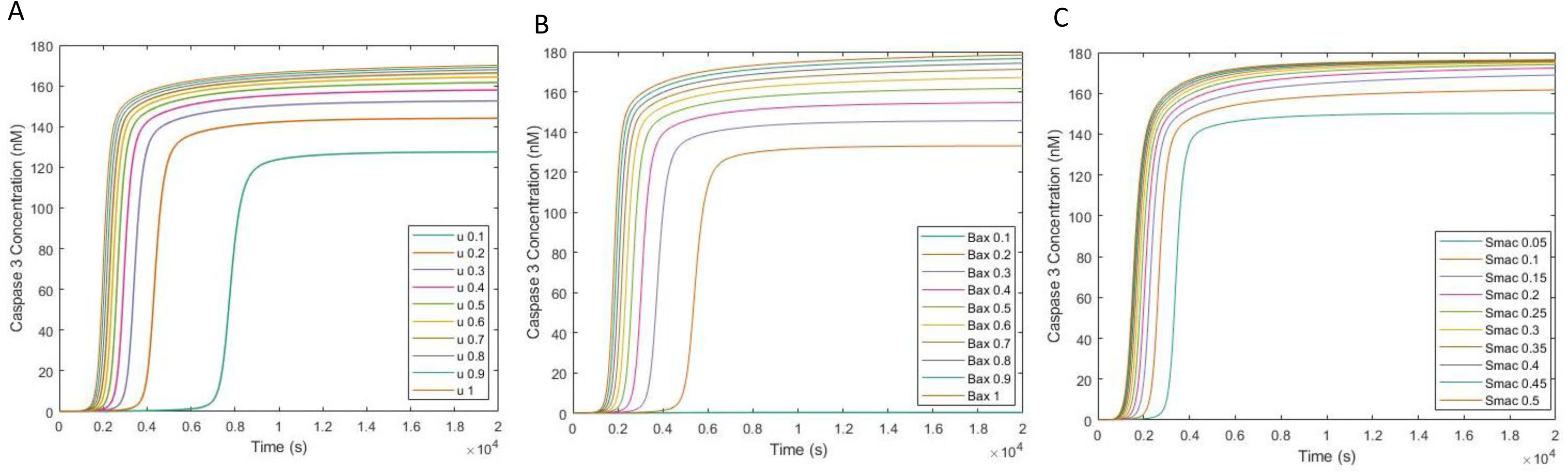
Impact of Input, BAX and SMAC concentration on Caspase 3 activation. Time course activation of caspase 3 upon step-like increase in A. Input stimulus concentration from 0.1-1nM, B. initial BAX concentration (BAX0) from 0.1-0.9nM and C. initial SMAC release rate from 0.05s^−1^- 0.5s^−1^. 4A. illustrates caspase 3 activation upon input stimulus variation. Lower *u* increases the caspase 3 activation time-delay while caspase 3 is activated earlier upon increase in *u*. 4B. indicates that caspase 3 simulated response time and amplitude is inversely related to initial BAX concentration (BAX0). Lowest BAX0 (0.1nM) results in caspase 3 activation time-delay of 500 seconds (∼8 mins) with an amplitude of over 130a.u., whereas the highest BAX0 (0.9nM) results in caspase 3 activation around 100 seconds (∼2 mins) with an amplitude greater than 160a.u. 4C. indicates that caspase 3 activation time-delay and amplitude is similarly inversely related to SMAC release rate. Lowest SMAC release rate (0.05s^−1^) results in an increase in caspase 3 activation time-delay to a lower amplitude, whereas the highest SMAC release rate (0.5s^−1^) results in earlier caspase 3 activation with a higher amplitude.

#### The model exhibits bistability

The Legewie model showed that IAPs were mediators of positive feedback causing bistability^20^. This and other models have shown that bistability is a key aspect in the regulation of biological networks and in particular is important for the determination of the irreversibility of apoptosis^20^. Therefore, we next wanted to investigate if the addition of BAX and SMAC to our model influenced the bistability and hysteresis previously demonstrated by the Legewie model. To confirm that the model exhibits bistable behaviour we performed a bifurcation analysis. Figure 3A shows two stable (solid line) and one unstable (dashed line) steady states. The system can reside in one of the two stable steady states (‘on’ or ‘off’) but not in the intermediate unstable steady state. In this case, the system remains at low caspase 3 activity (‘off’ state) for increasing stimuli until a threshold of stimulus concentration is reached, at which point caspase 3 activity switches to the second steady state (‘on’ state) irreversibly, in an all-or-none fashion. Critically, in line with time course simulations and literature findings^59,60^, the second bifurcation point of the system is negative(Figure 3B), which indicates that the activation of C3* is not reversible considering that it is not possible to have negative stimulus concentration. Depending on the state in which the system starts (‘on’ or ‘off’ state), differential response curves will be obtained^20^. Importantly, the first bifurcation point at *u*=0.06nM and the second at *u*=-0.01nM, indicate that the system displays hysteresis between stimulus concentrations of −0.01 to 0.06nM (Figure 3). This is in line with experimental observations which have shown that apoptosis activation depends on the intensity of the stimulus, allowing the cell to survive if the signal is not sufficient to drive the cell to begin programmed cell death^56^.

#### Bax and Smac regulate the timing of the apoptotic switch

The previous simulation demonstrated that the model that we have built recapitulates the *in vivo* behaviour of the intrinsic apoptosis pathway. Therefore, we were confident that this model could be used to study the influence of the pro-apoptotic regulators, BAX and SMAC on the mechanistic properties of the activation of the caspase cascade. Hence, we used simulations to investigate how BAX and SMAC control the timing of the apoptotic switch. We first investigated how the strength of the input stimulus (intensity of the signal) controls the observed time-lag of caspase activation (Figure 4A). A stimulus strength of u=0.5 nM demonstrated rapid C3* activation in an all-or-nothing fashion. Upon reduction of the input stimulus concentration, C3* does not activate until a later time point, demonstrating an increase in C3* activation time-delay, as is expected for a bistable system. Conversely, increasing the input stimulus concentration reduces the C3* activation time-delay (Figure 4A). Therefore, we observed that simulation response time is inversely related to the strength of the stimulus.

Next, we investigated whether alteration of BAX and SMAC activity could regulate this time-delay in C3* activity. Altering the initial concentration of BAX (0.1µM-1µM) or the release rate of SMAC (0.05s ^−1^ −0.5s ^−1^) results in a similar change in C3* time-delay. Higher concentrations of BAX and SMAC result in shorter C3* activation time-delay (Figures 4B and C). Together, these results show that the time-delay of C3* activation is controlled by (*i*) the strength of the stimulus, and (*ii*) the concentration of BAX and SMAC protein expression.

#### Bax and Smac regulate the threshold and amplitude of the apoptotic switch

The results so far indicate that the model that we have generated reflects the behaviour of the intrinsic apoptotic pathway and therefore can be used to analyse the relative contributions of BAX and SMAC to the irreversible activation of the system. Importantly, these proteins have been shown to be deregulated in several diseases such as cancer and neurodegeneration, where incorrect activation of apoptosis are hallmarks^4,6,26,28–42^. In the case of tumour cells, it has been shown that there is an increase of the threshold of apoptosis activation by downregulation of BAX and SMAC^16,31,35^. In neurodegeneration there is an improper activation of apoptosis that has been associated to increased levels of these regulatory proteins^39,40,61^. Thus, we posit our model could predict how alterations on BAX and SMAC levels can affect the intrinsic apoptosis pathway in these pathological conditions. To investigate at which precise values transition between stable steady states occur, indicating the threshold of caspase activation, we created bifurcation diagrams for differential concentrations of BAX and SMAC.

First, we investigated how changes in BAX concentration affect the system by performing bifurcation analysis with differential BAX levels. A range of bifurcation plots for differential BAX conditions (BAX0 = 0.1nM-1nM) is shown in Figure 5A. The first limit bifurcation point of the model with BAX0 at 0.5nM occurs at *u*=0.068nM, where the system transitions from the ‘off’ state to the ‘on’ state of C3* activity in an all-or-none fashion. With alteration in BAX0, differential system behaviour emerges. Upon decrease of BAX0, in a similar manner to what can be observed in cancer cells such as melanoma and colorectal cancer^6,28,34,36^, a higher limit bifurcation point is observed (u =0.6nM at BAX0=0.1nM). This shift in bifurcation point indicates an increase in threshold stimulus (*u*) required for activation of C3* activity upon decrease in BAX0. Conversely, with increase in BAX0 to 0.8nM, a decrease in threshold of C3* activation is observed with a reduced bifurcation point of u=0.04nM. This indicates that with a higher BAX0, there is a reduction in the required stimulus to reach the threshold of activation of C3* and can explain how cell death could be triggered at a lower drug concentration in pathologies were BAX levels are increased. Importantly, this is biologically sound considering that an increase in BAX translocation to the mitochondrial membrane will enhance MOMP, CytC release and reduce the stimulus required to activate the caspase cascade^5^.

**Figure 5.**
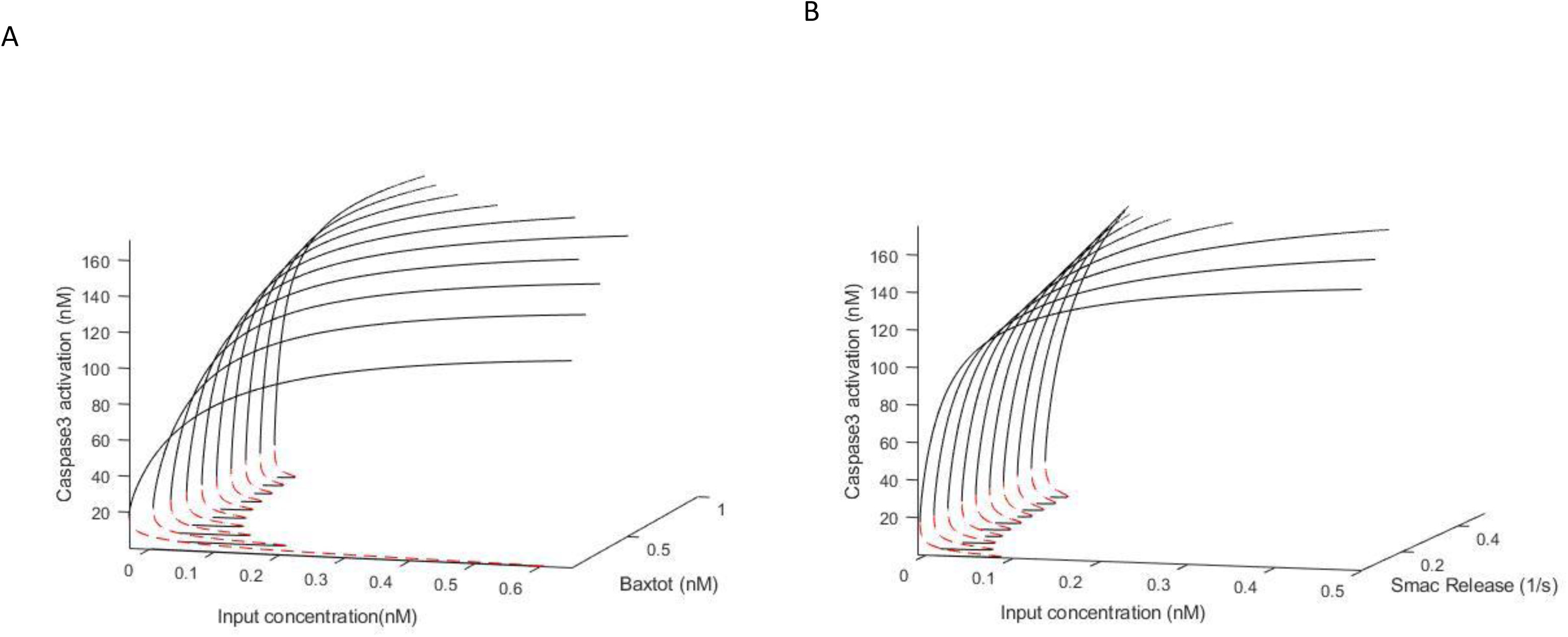
Impact of BAX and SMAC on the Activation Threshold of Caspase 3. (A). Bifurcation diagrams for a range of BAX initial concentrations (BAX0) (0.1-1nM). Nominal BAX0 is 0.5nM. The first limit bifurcation point of the nominal model with BAX0 at 0.5nM occurs at *u* (*0*.06 nM), where the system transitions from the ‘off’ state to the ‘on’ state of caspase 3 activity in an all-or-none fashion. Upon decrease of BAX0, a higher limit bifurcation point is observed (*u* (*0*.6nM) at BAX0=0.1nM). Conversely, with increase in BAX0 to 0.8nM, a decrease in threshold of caspase 3 activation is observed with a reduced bifurcation point of *u* (0.04nM). SMAC remained as in the nominal model for all simulations with SMAC release rate =0.1s^−1^. (B). Bifurcation diagrams for a range of SMAC release rate (0.05-0.5s-1). Nominal SMAC release rate is 0.1 s-1. The threshold for activation of C3* in the nominal model, with SMAC release rate (k19) = 0.1 is *u* (0.68nM). With decrease in SMAC activation by decreasing k19 to 0.01, the threshold for activation of Caspase 3 decreases to *u* (0.98nM) indicating the additional stimulus required to activate C3* upon decreased SMAC levels. With increase in SMAC activation (k19 = 0.1), a downward shift in activation threshold to *u* (0.02nM) is observed.

Next, we investigated how altered SMAC levels impact the transition between ‘off’ and ‘on’ C3* states. Figure 5B illustrates the C3* bifurcation plots for a range of SMAC values. The parameter ‘k19’ in Table 1 is the activation rate for SMAC and was altered over a range (0.05s^−1^ - 0.5s^−1^) to produce different levels of SMAC. BAX0 remained at 0.5nM throughout these simulations. The threshold for activation of C3* with k19=0.1s^−1^ is *u*=0.68nM). With decrease in SMAC activation by decreasing k19 to 0.05, the threshold for activation of caspase 3 increases to *u*=0.98nM indicating the additional stimulus required to activate C3* upon decreased SMAC levels. With increase in SMAC activation, a downward shift in C3* activation threshold to u=0.02nM is observed. This implies that with an increase in SMAC levels, by employing a faster activation rate (k19), a lower input stimulus is required to reach the threshold of activation of C3*.

These simulations demonstrate that our model can be used to predict what happens when alterations to SMAC and BAX levels occur in pathological conditions. Importantly, our simulations also showed that while alteration of BAX and SMAC impact the threshold of activation of C3* in a similar pattern, decrease of BAX0 causes a larger increase in the threshold of C3* activation when compared to a decrease in SMAC activation. This would indicate that BAX, but not SMAC, is a more prominent regulator of the apoptotic switch.

Next, we investigated how BAX and SMAC regulate the amplitude of caspase 3 activation. The bifurcation diagrams show BAX affected the C3* values of the ‘on’ state more prominently than the SMAC levels. Further, the ‘on’ state of caspase 3 saturates at high input concentrations (u>0.5nM). To analyse more quantitatively how alteration of BAX and SMAC impact this saturation amplitude of caspase 3 activation, we simulated dose responses for increasing input concentrations and different BAX and SMAC levels. Firstly, varying the BAX expression levels 5-fold down and up (0.1 – 1 nM) caused variations in the C3* amplitude from 120 to 180 nM (Supplementary Figure 1A). Note that for lower BAX level (<0.1) the activation threshold shifts to unphysiological high input concentrations and C3* does not activate. Secondly, varying the SMAC release rate over a 10-fold range (k19, 0.05 – 0.5 s^−1^) only caused variations in the C3* amplitude from 160 to 180 nM (Supplementary Figure 1B). Note that decreasing SMAC release rates even lower (<0.05s^−1^) C3* does not affect the ability of the system to switch on, nor does it change the C3* amplitudes. Thus, our model predicts that the amplitude of caspase 3 activation is more sensitive to changes in BAX than to changes in SMAC.

Finally, to get a further characterisation of the role of BAX and SMAC in the system we decided to investigate whether alteration of BAX or SMAC impacted the irreversibility of the switch. We studied the second bifurcation point of the system where C3* is transitioning from the high activity ‘on’ state to the ‘off’ state. We observed that alteration of BAX and SMAC does not impact the irreversibility of C3* activation. Although the second bifurcation point increased with increased BAX0 and SMAC, it never became positive indicating that the activation of C3* remains irreversible. Thus, BAX and SMAC do not impact the existence of bistability in our model. In line with other observations in the literature^21^, our results demonstrate that the main function of BAX and SMAC is to control the quantitative bistable properties (such as the “on” state threshold and amplitude), not to generate bistability itself. Bistability of our system emerges as a result of implicit and explicit feedback loops, as in line with the literature^19–21^.

#### Validation experiments confirm that caspase activation is more sensitive to BAX than SMAC

We performed experiments to validate the most salient model prediction; that alteration of BAX and SMAC expression levels impact the bistable properties of caspase activation. As outlined above, model simulations suggest that the amplitude of caspase activation can be impacted by alteration of BAX, and to a lesser extent SMAC, expression levels. To experimentally validate these model predictions and confirm the role that BAX and SMAC play in controlling the caspase 3 activation, we performed cell culture experiments. In these experiments, reduction of either BAX or SMAC level was achieved by RNA interference (siRNA) and intrinsic apoptosis was induced by the DNA damaging agent Etoposide (50µM for∼18 hours). To confirm that the downregulation of BAX and SMAC worked, their protein expression levels were measured by western blot (Figure 6A). Western blot also demonstrated that upon Etoposide treatment, an increase in BAX protein levels are observed, independently of SMAC levels. This increase in BAX protein levels indicate the induction of BAX upon DNA damage to translocate to the mitochondria and initiate the intrinsic caspase cascade.

**Figure 6.**
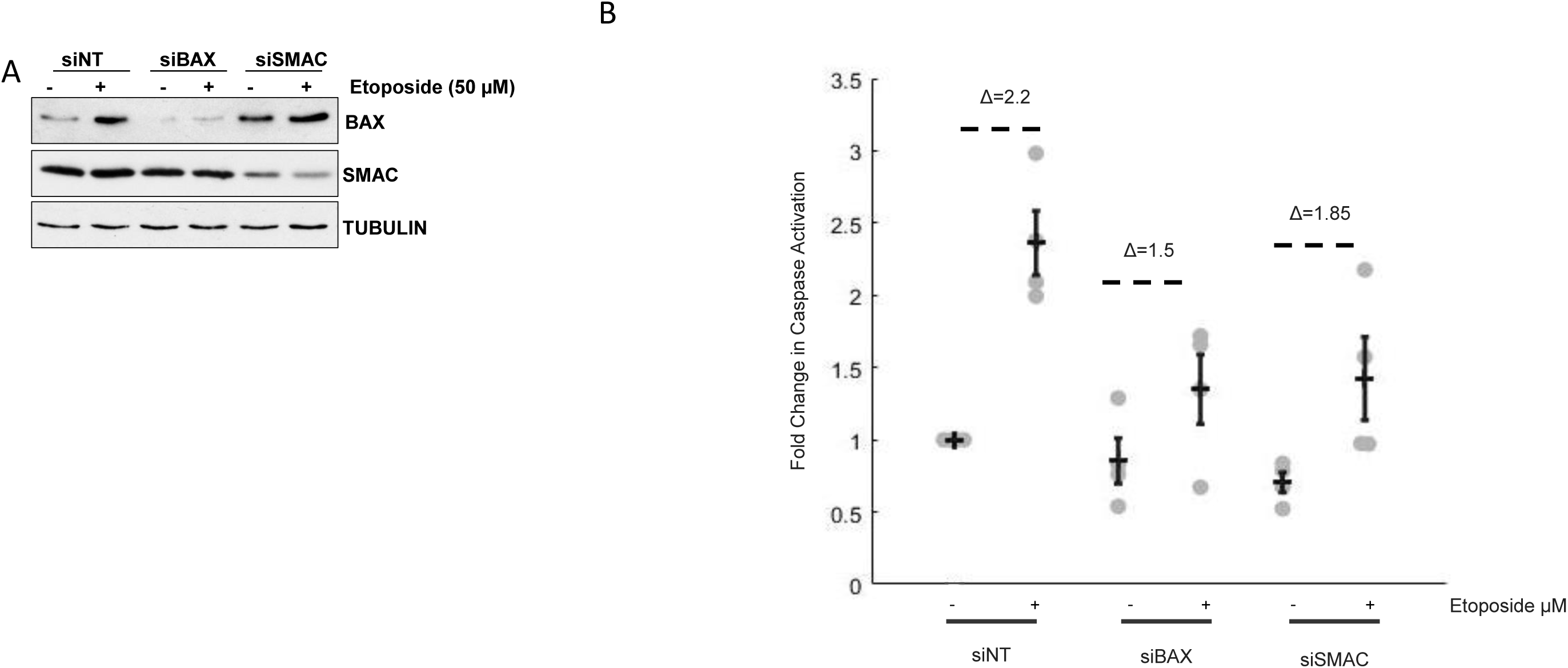
siRNA-mediated downregulation of BAX and SMAC reduces caspase activation upon Etoposide treatment. (A) HeLa cells were transfected with either BAX, SMAC or non-target (NT) siRNA and 36 hours after transfection the cells were treated with 50 µM Etoposide for 18 hours in growing conditions where indicated. Cell lysates were collected, and expression levels of BAX and SMAC proteins were monitored by blotting with specific antibodies. Tubulin α was used as loading control. B) HeLa cells treated as in A were collected from the media or after trypsinization and caspase activation was assessed by FITC-VAD-FMK binding and measured by flow cytometry. Fold change in percentage of FITC-VAD-FMK positive cells is represented and normalized against control cells (siRNA non target-transfected cells treated with DMSO) as annotated by Δ. Error bars show SD=0, 0.59, 0.31, 0.48, 0.14, 0.58 for siNT DMSO, siNT Etoposide, siBAX DMSO, siBAX Etoposide, siSMAC DMSO and siSMAC Etoposide groups respectively. N=4.

The corresponding total caspase activity was measured by flow cytometry and is shown in Figure 6B. Non-transfected control cells show a basal caspase activity fold change of 2.2 upon application of Etoposide. Upon siRNA-mediated down regulation of BAX, there is no change in basal caspase activation levels. Treatment with etoposide induced a 1.5 fold increase in caspase activation, constituting a 0.7 fold difference to vehicle treated control cells. Upon SMAC downregulation, the onset of basal caspase activation in vehicle treated cells is reduced. Upon Etoposide treatment, an increase in caspase activation by 1.85 fold was observed, a 0.35 fold difference to vehicle treated control cells. These experimental results confirm the model predictions that upon downregulation of BAX and SMAC, a decreased amplitude in caspase activation is observed and that BAX downregulation has a more pronounced impact than SMAC downregulation.

#### Simulated Caspase Activation Correlates with Drug Sensitivity in a Panel of Melanoma Cell Lines

To further experimentally validate our model, we next tested whether our model could explain the drug-responses of different cell lines. To do this, we analysed publicly available NCI-60 datasets for which proteomic-profiling and drug-response data were available and used the protein expression information to perform cell line specific simulations. These datasets consist of 60 cell lines from 9 tissues, mass-spectrometry based proteome measurements, and measured drug response scores for more than 100,000 drugs, including etoposide^62,63^. BAX but not SMAC levels correlated with etoposide response, Pearson correlation coefficient (C) = 0.307, p-value (p) =0.019, (Suppl Fig. 3), suggesting that BAX but not SMAC protein levels provide useful information to personalize the model to the individual cell lines. Thus, we focused on BAX to generate cell-line specific simulations. The measured SWATH values for BAX were used to adjust the BAX total protein expression parameter in the model as follows; if for example BAX levels were 1.5 times increased in cell line A, then we increased the corresponding BAX protein expression parameter (k1) by a factor of 1.5. The resulting set of cell-line specific parameters was then used to simulate the caspase activation trajectories for each cell line.

The simulation results showed that the C3 amplitude and activation-threshold varied markedly between cell-lines (ranging from 120 to almost 180 nM, Fig 7A, B) and correlated with the etoposide response (C=0.3, p-value=0.022, Fig 7C). This correlation between the C3 amplitude and the etoposide response might be a tissue effect, meaning that one tissue is sensitive to etoposide, but another is not. To check this, we looked at the variability of the simulated responses across tissues but observed no pronounced differences of the C3 amplitude between different tissues (Fig. 7A). This indicates that the observed correlation between the C3 amplitude and the etoposide response depends on the specific genomics of each cell-line and not tissue-specific differences.

**Figure 7.**
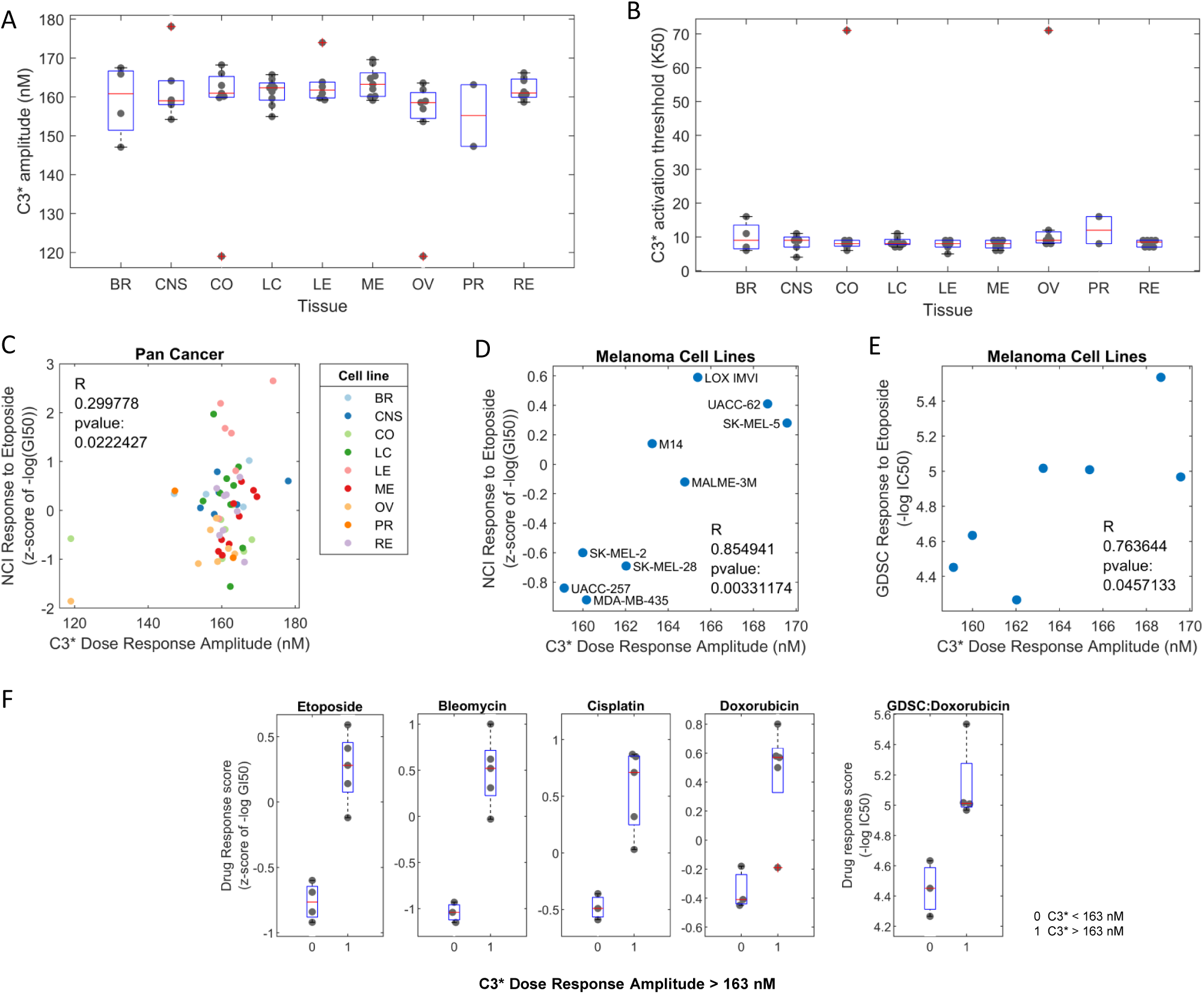
Cell Line Specific Simulations Correlate with Drug Response. (A,B) Distributions of the cell-line specific simulated caspase 3 activation amplitudes (A) and caspase 3 activation thresholds (B) for the issues indicated on the x-axis. Boxplots indicate the upper and lower quartile (blue box) and median (red line). Dots represent the simulated values for each cell line. BR: breast, CNS: central nervous system, CO: colorectal, LC: lung cancer, LE: leukaemia; ME: melanoma, OV: ovarian cancer, PR: prostate, RE: renal cancer. (C) Pan cancer cell line specific simulations were carried out and C3 amplitude was correlated with cell line response to Etoposide (C=0.3, pval=0.022). (D) Melanoma cell line specific simulations(n=9) indicated a positive correlation with response to etoposide measured by the NCI-60 screen (C=0.855, pval = 0.003). (E) Melanoma cell line specific simulations (n=7) are positively correlated with etoposide response measured in the GDSC screen (C =0.76, pval=0.046). (F) Classification of the melanoma cells into sensitive and insensitive cell lines. Shown are the distributions of the indicated drug responses (y-axis) for cells with a low simulated caspase 3 activation amplitude (<163 nM) on the left (0) and cells with a high amplitude (>163 nM) on the right (1). Boxplots indicate the upper and lower quartile (blue box) and median (red line). Dots represent the values for each cell line.

Another explanation for the correlation might be that model explains the etoposide response very well in one tissue but not the others. To test this, we analysed the simulations for each tissue separately, focusing on those tissues with more than 5 cell lines (Table 3). The best correlation between the simulated C3 amplitude and the etoposide response was observed in melanoma cell lines, C=0.855, pval =0.003, n=9 cell-lines (Fig 7D). To confirm this, we also analysed data from an independent drug screen (GDSC^64^), and found a similar correlation C=0.76, pval0.046, n=7 (Fig 7E). Ovarian cancer cell lines also exhibited a correlation, C=0.77, pval =,0.04, n=7 (Fig S3). The correlations for the other tissues (leukemia, colorectal, CNS, breast, renal, lung and prostate) were weak and not significant (Table 3).

**Table 3.** Table of Cell Lines used for Cell Line Specific Simulations.

| Tissue | Correlation (C) | P-Value |
| --- | --- | --- |
| Leukemia | 0.575 | 0.233 |
| Colorectal | 0.046 | 0.923 |
| CNS | 0.428 | 0.397 |
| Breast | 0.352 | 0.648 |
| Renal | -0.237 | 0.572 |
| Lung | -0.362 | 0.339 |
| Ovarian | 0.759 | 0.048 |
| Melanoma | 0.845 | 0.004 |

Focusing on the melanoma cells, we wanted to test how general the model is with respect to other DNA damaging drugs and analysed the data for bleomycin, cisplatin, and doxorubicin. For all three drugs, the drug response scores correlated with the simulated C3 amplitude (Fig S3). The correlation of the C3 amplitude with the bleomycin and doxorubicin responses was particularly strong and significant (C=0.77, pval=0.03 and C=0.79, pval=0.02, respectively). The results suggest that the model can be used to simulate and predict caspase activation and the responses to DNA damaging drugs in melanoma cells.

Given the observed correlation between the simulated C3 amplitude and the response to DNA damaging drugs, we asked whether a particular threshold in the model could distinguish between sensitive and resistant cells. We found that for all analysed drugs, a threshold value for the C3 amplitude of 163 nM was able to distinguish between sensitive and resistant cells (Fig. 7F). All melanoma cells with a simulated C3 amplitude less than 163 nM were sensitive to etoposide, bleomycin, cisplatin and doxorubicin, with NCI-60 response scores < 0 (n=8 cells). Conversely, the cell lines with a simulated C3 amplitude greater that 163 nM were resistant to all three drugs. Only one outlier for doxorubicin was observed, where M14, was falsely classified as sensitive (Fig. 7F). In summary, our results suggest that BAX protein levels can be used to personalize the model, simulate caspase activation, and predict drug-responses for DNA-damaging drugs in melanoma cells. A defined threshold value of 163 nM of caspase activation in the model distinguishes between sensitive and resistant melanoma cells.

## Discussion

In the current work we have developed a mathematical model of the intrinsic apoptotic pathway that recapitulates the behaviour of this systems in cells. The current model includes BAX and SMAC, two nodes that have not been widely analysed in previous computational models but are *bona fide* regulators of this network. Importantly, the addition of these nodes in the model and their experimental is a clear advance of our study with respect to other models, and better recapitulates the behaviour of the system in cells and the enhances the predictive power of the computational model. The most salient model prediction was that BAX exerts stronger control over the amplitude of the apoptotic switch than SMAC, and our cell culture experiments validated this prediction. Using measured BAX protein levels to personalise the model to each cell line of the NCI-60 panel, it was revealed that the simulated caspase activity correlated with the cell line etoposide response. Further, a distinct threshold of caspase 3 activation separated sensitive from insensitive melanoma lines for a variety of DNA damaging drugs. This finding is particularly interesting in cancer and provides a possible explanation for why the deregulation of BAX is more common than deregulation of SMAC^5,65,66^.

Importantly, our model confirms the existence of bistability in accordance with other studies of the apoptosis network^20,21^. However, many studies modelling apoptosis focused on the extrinsic pathway consisting of death receptor activation, subsequent activation of caspases and secondary activation of the mitochondrial pathway^22,23,67,68^, or the processes upstream of caspase activation such as BCL2 protein regulation of MOMP^21–23,69,70^. Other modelling studies that concerned the activation of caspases in the intrinsic pathway did either not look at the control of bistable properties^22,67,69^, or did not include BAX and SMAC^19,20,23^. A notable exception is the work by Tyson and colleagues^21^, which provided and excellent account of which network nodes and parameters regulate the bistability properties, in particular irreversibility, time-delay and threshold. They found that the time-delay and threshold are primarily controlled by the initiator module (corresponding to BAX in our model), which is in accordance with our findings. However, we also find that SMAC, which merely relayed the signal in the Tyson model, markedly controlled the delay and threshold of caspase activation. Further, and in contrast to Tyson, we also analysed how BAX and SMAC control the amplitude of the apoptotic switch, and experimentally validated this prediction.

The results from our simulations clearly indicate that the modification of BAX and SMAC levels have a direct effect in the threshold and time delay of caspase 3 activation and determine when the system reaches the point of no return. Thus, the simulation shows that decrease of both proteins results in an increase of this threshold of apoptosis, i.e. lowering the expression of these proteins makes the system more resistant to pro-apoptotic stimuli. Therefore, at low level of expression of BAX, our model mimics the cellular state of tumours where the loss of BAX expression is a common feature. Conversely, high level of expression BAX would resemble other diseases were aberrant activation of apoptosis is part of the pathology. For instance, increases in BAX level have been associated with the activation of programmed cell death in degenerative diseases, especially neurodegeneration, and other diseases such as ischemia and heart infarction^71,72^. With respect to SMAC, similar observations are shown in the same diseases^4,38,39,41,50,73–81^. For these reasons, our bifurcation analysis can be considered as simulations of what would happen to the whole system in physiological and pathological conditions. The fact that the personalised simulations of caspase 3 activity correlated with the response to DNA damaging drugs in the NCI-60 pan cancer dataset suggests that the model can be used to predict the individual cellular responses of different cell lines and patients. This could work particularly well for melanoma, where the cell-line specific simulations confirmed the existence of a distinct threshold separating sensitive from resistant melanoma cells. To improve the model for other tissues, further key nodes that regulate the caspase system in addition to BAX could be tested, mathematically modelled, and connected to the current model. Prime candidates are members of the BCL2 family and a more detailed model of the outer mitochondria membrane permeabilization that have proven predictive power in triple negative breast and colorectal cancer ^44,82,83^. Personalising this model with experimental and clinical data could be used to carry out *in silico* drug targeting and could help the development of novel therapeutics for apoptosis-dependent diseases. Importantly, our simulations indicate that critical bistable properties such as the amplitude of the “on-state” are more dependent on variations of the level of BAX than SMAC. Taking into account that there are currently different SMAC and BAX mimetics for the treatment of cancer at different stages of development, our model would support the idea that BAX mimetics would be more effective than SMAC mimetics. In the context of regenerative therapies that aim to preventing excessive apoptosis, our finding might indicate that targeting BAX translocation could be more efficient than targeting SMAC.

In summary we present a new experimentally validated mathematical model that can be used to study the dynamics of activation of apoptosis by the mitochondrial pathway.

## Materials and Methods

### Model Construction and Simulation

Our core model is based on a well-established model of caspase bistability in the literature ^20^ and has been extended to include BAX and SMAC and the associated reactions (see main text, Fig. 1). Based on this reaction kinetic scheme an ODE model has been constructed using the principle of mass balance and the appropriate reaction kinetic laws, as is common practice^84^. All ODE simulations were carried out using the MATLAB environment (R2018a). Bifurcation analyses were carried out using XPPAUT (http://www.math.pitt.edu/~bard/bardware/xpp/xpp.html) and plotted using the plotxppaut plugin for MATLAB(http://www2.gsu.edu/~matrhc/XPP-Matlab.html).

### Cell culture

HeLa cells (ATTC) were grown in DMEM (Gibco) supplemented with 10% FBS (Gibco) and 2 mM glutamine (Gibco) at 37°C, 21% O_2_ and 5% CO_2_ in a humidified incubator and kept in exponentially growing conditions. Lipofectamine 2000 (Invitrogen) was used for siRNA transfection according to manufacturer’s protocol. Briefly, cells were seeded 24 hours prior to transfection (70% confluency) and 6.25 µL of Lipofectamine 2000 and 100 pmoles of siRNA per 60 mm petri dish were used. 24 hours after transfection, cells were treated with 50 µM of Etoposide (Sigma Aldrich) or vehicle (DMSO) for 18 hours. siRNAs against human BAX and SMAC and a non-target siRNA control were purchased from Dharmacon (ON-TARGETplus Human BAX (581) siRNA – SMARTpool, ON-TARGETplus Human DIABLO (56616) siRNA – SMARTpool and ON-TARGETplus Non-targeting siRNA #1, respectively)

### Caspase activation assay

Caspase activation was measured in an Accuri’s C6 Flow Cytometer System using the CaspACE-FITC-VAD-FMK kit (Promega). Alive and dead cells were collected, centrifuged for 5 minutes at 300x*g* and incubated with FITC-VAV-FMK in serum free media (1:2000) for 30 minutes at 37°C protected from light. Cells were washed once with PBS and incubated with PI for 5 minutes at room temperature prior to acquisition. FITC and PI emissions were measured using the 533/30 and 670LP detectors respectively following excitation by 488nm laser. Statistical analysis was carried out to determine the mean caspase activation and standard deviation within each group (siNT, siBAX and siSMAC), control and treated. Fold change in caspase activation was calculated for each treated group compared to the group control.

### Western Blot

Cells were lysed in a buffer containing: 20 mM Hepes pH 7.5, 150 mM NaCl, 1 % NP40 supplemented with protease and phosphatase inhibitors (Roche). Protein extracts were clarified by centrifugation (14,000 rpm, 4°C, 10 minutes) and pellet discarded. Protein quantification was carried out using the BCA protein Assay kit (23225-Invitrogen) according to manufacturer’s protocol. Primary antibodies (1:1000 dilution): anti-SMAC Y12 (ab32023-Abcam), anti-BAX 6A7 (556467-BD Pharmingen), Tubulin TU-U2 (sc-8035-Santa Cruz).

### Cell Line Specific Simulations

BAX and SMAC protein expression (SWATH values^85,86^), GDSC Etoposide response and NCI-60 drug response to Etoposide, Doxorubicin, Bleomycin and Cisplatin were downloaded using the CellMiner tool (https://discover.nci.nih.gov/cellminercdb/, (29/11/2019). All information was collated and stored in a single file(Access). The measured SWATH values of the BAX protein levels swaBAX (log10) were mean normalised *relBAX*_*i*_ = *swaBAX*_*i*_ − *mean*(*swaBAX*_*i*_) used to adjust the *BAX*_*tot*_ parameter for each cell line *i* as follows 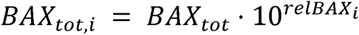, with the nominal BAX protein expression parameter of *BAX*_*tot*_ = 0.5.

### Data availability

The MATLAB and XPPAUT files and relevant NCI-60 data of the model are available in the supplement. SBML version of the code is available at EMBL-EBI BioModels (accession here when obtained).

## Acknowledgements

Authors wold like to acknowledge the UCD Conway Core Flow Cytometry Facility and Dr Alfonso Blanco for flow cytometry support. This work was supported by the Science Foundation Ireland Grant 15_CDA_3495.

## Author Contributions

All Authors contributed to writing the manuscript. SMK, DF, DM conceived the study and analysed the data. SMK, DF constructed the model and SMK carried out the simulations. LG and SMK carried out the experimental work.

## References

1. Chung C. Restoring the switch for cancer cell death: Targeting the apoptosis signaling pathway. Am J Heal Pharm. 2018;75(13):945–952. doi:10.2146/ajhp170607

2. Elmore S. Apoptosis: a review of programmed cell death. Toxicol Pathol. 2007;35(4):495–516. doi:10.1080/01926230701320337

3. Cohen GM. Caspases: the executioners of apoptosis. Biochem J. 1997;326 (Pt 1):1–16. http://www.ncbi.nlm.nih.gov/pubmed/9337844. Accessed December 5, 2018.

4. Maes ME, Schlamp CL, Nickells RW. BAX to basics: How the BCL2 gene family controls the death of retinal ganglion cells. Prog Retin Eye Res. 2017;57:1–25. doi:10.1016/J.PRETEYERES.2017.01.002

5. Kalkavan H, Green DR. MOMP, cell suicide as a BCL-2 family business. Cell Death Differ. 2018;25(1):46–55. doi:10.1038/cdd.2017.179

6. Anvekar RA, Asciolla JJ, Missert DJ, Chipuk JE. Born to be alive: a role for the BCL-2 family in melanoma tumor cell survival, apoptosis, and treatment. Front Oncol. 2011;1(34). doi:10.3389/fonc.2011.00034

7. Finucane DM, Bossy-Wetzel E, Waterhouse NJ, Cotter TG, Green DR. Bax-induced caspase activation and apoptosis via cytochrome c release from mitochondria is inhibitable by Bcl-xL. J Biol Chem. 1999;274(4):2225–2233. doi:10.1074/JBC.274.4.2225

8. Berthelet J, Dubrez L. Regulation of Apoptosis by Inhibitors of Apoptosis (IAPs). Cells. 2013;2(1):163–187. doi:10.3390/cells2010163

9. Galbán S, Duckett CS. XIAP as a ubiquitin ligase in cellular signaling. Cell Death Differ. 2010;17(1):54–60. doi:10.1038/cdd.2009.81

10. Salvesen GS, Duckett CS. IAP proteins: Blocking the road to death’s door. Nat Rev Mol Cell Biol. 2002;3(6):401–410. doi:10.1038/nrm830

11. Hill MM, Adrain C, Duriez PJ, Creagh EM, Martin SJ. Analysis of the composition, assembly kinetics and activity of native Apaf-1 apoptosomes. EMBO J. 2004;23(10):2134–2145. doi:10.1038/sj.emboj.7600210

12. Du C, Fang M, Li Y, Li L, Wang X. Smac, a mitochondrial protein that promotes cytochrome c-dependent caspase activation by eliminating IAP inhibition. Cell. 2000;102(1):33–42. http://www.ncbi.nlm.nih.gov/pubmed/10929711. Accessed May 15, 2019.

13. Chai J, Du C, Wu J-W, Kyin S, Wang X, Shi Y. Structural and biochemical basis of apoptotic activation by Smac / DIABLO. doi:10.1038/35022514

14. Rehm M, Düßmann H, Prehn JHM. Real-time single cell analysis of Smac/DIABLO release during apoptosis. J Cell Biol. 2003;0913(6):21–9525. doi:10.1083/jcb.200303123

15. Kocab AJ, Duckett CS. Inhibitor of apoptosis proteins as intracellular signaling intermediates. FEBS J. 2016;283(2):221–231. doi:10.1111/FEBS.13554

16. Chung C. Restoring the switch for cancer cell death: Targeting the apoptosis signaling pathway. Am J Health Syst Pharm. 2018;75(13):945–952. doi:10.2146/ajhp170607

17. Fallahi E, O’Driscoll N, Matallanas D. The MST/Hippo Pathway and Cell Death: A Non-Canonical Affair. Genes (Basel). 2016;7(6):28. doi:10.3390/genes7060028

18. Fussenegger M, Bailey JE, Varner J. A mathematical model of caspase function in apoptosis. Nat Biotechnol. 2000;18(7):768–774. doi:10.1038/77589

19. Eissing T, Conzelmann H, Gilles ED, Allgöwer F, Bullinger E, Scheurich P. Bistability analyses of a caspase activation model for receptor-induced apoptosis. J Biol Chem. 2004;279(35):36892–36897. doi:10.1074/jbc.M404893200

20. Legewie S, Blüthgen N, Herzel H. Mathematical Modeling Identifies Inhibitors of Apoptosis as Mediators of Positive Feedback and Bistability. PLoS Comput Biol. 2006;2(9):e120. doi:10.1371/journal.pcbi.0020120

21. Zhang T, Brazhnik P, Tyson JJ. Computational Analysis of Dynamical Responses to the Intrinsic Pathway of Programmed Cell Death. Biophys J. 2009;97(2):415–434. doi:10.1016/j.bpj.2009.04.053

22. Albeck JG, Burke JM, Spencer SL, Lauffenburger DA, Sorger PK. Modeling a snap-action, variable-delay switch controlling extrinsic cell death. Levchenko A, ed. PLoS Biol. 2008;6(12):2831–2852. doi:10.1371/journal.pbio.0060299

23. Bagci EZ, Vodovotz Y, Billiar TR, Ermentrout GB, Bahar I. Bistability in Apoptosis: Roles of Bax, Bcl-2, and Mitochondrial Permeability Transition Pores. Biophys J. 2006;90(5):1546–1559. doi:10.1529/BIOPHYSJ.105.068122

24. Ballweg R, Paek AL, Zhang T. A dynamical framework for complex fractional killing. Sci Rep. 2017;7(1):8002. doi:10.1038/s41598-017-07422-2

25. Chang LK, Putcha G V, Deshmukh M, Johnson EM. Mitochondrial involvement in the point of no return in neuronal apoptosis. Biochimie. 84(2-3):223–231. http://www.ncbi.nlm.nih.gov/pubmed/12022953. Accessed December 5, 2018.

26. Kholoussi NM, El-Nabi SEH, Esmaiel NN, Abd El-Bary NM, El-Kased AF. Evaluation of Bax and Bak gene mutations and expression in breast cancer. Biomed Res Int. 2014;2014:249372. doi:10.1155/2014/249372

27. Moshynska O, Sankaran K, Saxena A. Molecular detection of the G(−248)A BAX promoter nucleotide change in B cell chronic lymphocytic leukaemia. Mol Pathol. 2003;56(4):205–209. http://www.ncbi.nlm.nih.gov/pubmed/12890741. Accessed March 25, 2019.

28. Tai YT, Lee S, Niloff E, Weisman C, Strobel T, Cannistra SA. BAX protein expression and clinical outcome in epithelial ovarian cancer. J Clin Oncol. 1998;16(8):2583–2590. doi:10.1200/JCO.1998.16.8.2583

29. Qin S, Yang C, Li S, Xu C, Zhao Y, Ren H. Smac: Its role in apoptosis induction and use in lung cancer diagnosis and treatment. Cancer Lett. 2012;318(1):9–13. doi:10.1016/J.CANLET.2011.12.024

30. Favaloro B, Allocati N, Graziano V, Di Ilio C, De Laurenzi V. Role of apoptosis in disease. Aging (Albany NY). 2012;4(5):330–349. doi:10.18632/aging.100459

31. Fulda S. Inhibitor of apoptosis proteins in pediatric leukemia: molecular pathways and novel approaches to therapy. Front Oncol. 2014;4:3. doi:10.3389/fonc.2014.00003

32. Fulda S. Promises and Challenges of Smac Mimetics as Cancer Therapeutics. Clin Cancer Res. 2015;21(22):5030–5036. doi:10.1158/1078-0432.CCR-15-0365

33. Wang YF, Jiang CC, Kiejda KA, Gillespie S, Zhang XD, Hersey P. Apoptosis induction in human melanoma cells by inhibition of MEK is caspase-independent and mediated by the Bcl-2 family members PUMA, Bim, and Mcl-1. Clin Cancer Res. 2007;13(16):4934–4942. doi:10.1158/1078-0432.CCR-07-0665

34. Krajewski S, Blomqvist C, Franssila K, et al. Reduced expression of proapoptotic gene BAX is associated with poor response rates to combination chemotherapy and shorter survival in women with metastatic breast adenocarcinoma. Cancer Res. 1995;55(19):4471–4478. http://www.ncbi.nlm.nih.gov/pubmed/7671262. Accessed April 23, 2019.

35. Woo Lee J, Goo Jeong E, Hwa Soung Y, et al. Decreased expression of tumour suppressor Bax-interacting factor-1 (Bif-1), a Bax activator, in gastric carcinomas. Pathology. 2006;38(4):312–315. doi:10.1080/00313020600820880

36. Jansson A, Sun X-F. *Bax* Expression Decreases Significantly From Primary Tumor to Metastasis in Colorectal Cancer. J Clin Oncol. 2002;20(3):811–816. doi:10.1200/JCO.2002.20.3.811

37. Liu Z, Ding Y, Ye N, Wild C, Chen H, Zhou J. Direct Activation of Bax Protein for Cancer Therapy. Med Res Rev. 2016;36(2):313–341. doi:10.1002/med.21379

38. Kermer P, Liman J, Weishaupt JH, Bähr M. Neuronal Apoptosis in Neurodegenerative Diseases: From Basic Research to Clinical Application. Rev Neurodegener Dis. 2004;1:9–19. doi:10.1159/000076665

39. Vila M, Jackson-Lewis V, Vukosavic S, et al. Bax ablation prevents dopaminergic neurodegeneration in the 1-methyl-4-phenyl-1,2,3,6-tetrahydropyridine mouse model of Parkinson’s disease. Proc Natl Acad Sci. 2001;98(5):2837–2842. doi:10.1073/pnas.051633998

40. Lu G, Kwong W, Li Q, Wang X, Feng Z, Yew D. bcl2, bax, and nestin in the brains of patients with neurodegeneration and those of normal aging. J Mol Neurosci. 2005;27(2):167–174. doi:10.1385/JMN:27:2:167

41. Perier C, Tieu K, Guegan C, et al. Complex I deficiency primes Bax-dependent neuronal apoptosis through mitochondrial oxidative damage. Proc Natl Acad Sci. 2005;102(52):19126–19131. doi:10.1073/pnas.0508215102

42. Montero J, Letai A. Why do BCL-2 inhibitors work and where should we use them in the clinic? Cell Death Differ. 2018;25(1):56–64. doi:10.1038/cdd.2017.183

43. Hong J-Y, Kim G-H, Kim J-W, et al. Computational modeling of apoptotic signaling pathways induced by cisplatin. doi:10.1186/1752-0509-6-122

44. Lindner AU, Concannon CG, Boukes GJ, et al. Systems Analysis of BCL2 Protein Family Interactions Establishes a Model to Predict Responses to Chemotherapy. Cancer Res. 2013;73(2):519–528. doi:10.1158/0008-5472.CAN-12-2269

45. Flanagan L, Sebastia J, Delgado ME, Lennon JC, Rehm M. Dimerization of Smac is crucial for its mitochondrial retention by XIAP subsequent to mitochondrial outer membrane permeabilization. Biochim Biophys Acta - Mol Cell Res. 2011;1813(5):819–826. doi:10.1016/J.BBAMCR.2011.02.011

46. Liu Z, Sun C, Olejniczak ET, et al. Structural basis for binding of Smac/DIABLO to the XIAP BIR3 domain. Nature. 2000;408(6815):1004–1008. doi:10.1038/35050006

47. Zou H, Yang R, Hao J, et al. Regulation of the Apaf-1/caspase-9 apoptosome by caspase-3 and XIAP. J Biol Chem. 2003;278(10):8091–8098. doi:10.1074/jbc.M204783200

48. Bratton SB, Walker G, Srinivasula SM, et al. Recruitment, activation and retention of caspases-9 and −3 by Apaf-1 apoptosome and associated XIAP complexes. EMBO J. 2001;20(5):998–1009. doi:10.1093/emboj/20.5.998

49. Qin S, Yang C, Li S, Xu C, Zhao Y, Ren H. Smac: Its role in apoptosis induction and use in lung cancer diagnosis and treatment. Cancer Lett. 2012;318(1):9–13. doi:10.1016/J.CANLET.2011.12.024

50. Du C, Fang M, Li Y, Li L, Wang X. Smac, a Mitochondrial Protein that Promotes Cytochrome c–Dependent Caspase Activation by Eliminating IAP Inhibition. Cell. 2000;102(1):33–42. doi:10.1016/S0092-8674(00)00008-8

51. Fletcher JI, Huang DCS. Controlling the cell death mediators Bax and Bak: puzzles and conundrums. Cell Cycle. 2008;7(1):39–44. doi:10.4161/cc.7.1.5178

52. Westphal D, Dewson G, Czabotar PE, Kluck RM. Molecular biology of Bax and Bak activation and action. Biochim Biophys Acta. 2011;1813(4):521–531. doi:10.1016/j.bbamcr.2010.12.019

53. MacFarlane M, Merrison W, Bratton SB, Cohen GM. Proteasome-mediated degradation of Smac during apoptosis: XIAP promotes Smac ubiquitination in vitro. J Biol Chem. 2002;277(39):36611–36616. doi:10.1074/jbc.M200317200

54. Ali R, Singh S, Haq W. IAP Proteins Antagonist: An Introduction and Chemistry of Smac Mimetics under Clinical Development. Curr Med Chem. 2018;25(31):3768–3795. doi:10.2174/0929867325666180313112229

55. Taylor RC, Cullen SP, Martin SJ. Apoptosis: controlled demolition at the cellular level. Nat Rev Mol Cell Biol. 2008;9(3):231–241. doi:10.1038/nrm2312

56. Tang HL, Tang HM, Mak KH, et al. Cell survival, DNA damage, and oncogenic transformation after a transient and reversible apoptotic response. Luo K, ed. Mol Biol Cell. 2012;23(12):2240–2252. doi:10.1091/mbc.e11-11-0926

57. Takemoto K, Nagai T, Miyawaki A, Miura M. Spatio-temporal activation of caspase revealed by indicator that is insensitive to environmental effects. J Cell Biol. 2003;160(2):235–243. doi:10.1083/JCB.200207111

58. Widmann C, Gibson S, Johnson GL. Caspase-dependent cleavage of signaling proteins during apoptosis. A turn-off mechanism for anti-apoptotic signals. J Biol Chem. 1998;273(12):7141–7147. doi:10.1074/JBC.273.12.7141

59. Chang LK, Putcha GV, Deshmukh M, Johnson EM. Mitochondrial involvement in the point of no return in neuronal apoptosis. Biochimie. 2002;84(2-3):223–231. doi:10.1016/S0300-9084(02)01372-X

60. Deshmukh M, Kuida K, Johnson EM. Caspase Inhibition Extends the Commitment to Neuronal Death Beyond Cytochrome c Release to the Point of Mitochondrial Depolarization. J Cell Biol. 2000;150(1):131–144. doi:10.1083/JCB.150.1.131

61. Hsu M-J, Sheu J-R, Lin C-H, Shen M-Y, Hsu CY. Mitochondrial mechanisms in amyloid beta peptide-induced cerebrovascular degeneration. Biochim Biophys Acta - Gen Subj. 2010;1800(3):290–296. doi:10.1016/J.BBAGEN.2009.08.003

62. Reinhold WC, Sunshine M, Liu H, et al. CellMiner: A web-based suite of genomic and pharmacologic tools to explore transcript and drug patterns in the NCI-60 cell line set. Cancer Res. 2012;72(14):3499–3511. doi:10.1158/0008-5472.CAN-12-1370

63. Shankavaram UT, Varma S, Kane D, et al. CellMiner: a relational database and query tool for the NCI-60 cancer cell lines. BMC Genomics. 2009;10(1):277. doi:10.1186/1471-2164-10-277

64. Yang W, Soares J, Greninger P, et al. Genomics of Drug Sensitivity in Cancer (GDSC): a resource for therapeutic biomarker discovery in cancer cells. Nucleic Acids Res. 2013;41(Database issue):D955–61. doi:10.1093/nar/gks1111

65. Hassan M, Watari H, AbuAlmaaty A, Ohba Y, Sakuragi N. Apoptosis and molecular targeting therapy in cancer. Biomed Res Int. 2014;2014:150845. doi:10.1155/2014/150845

66. Amundson SA, Myers TG, Scudiero D, Kitada S, Reed JC, Fornace AJ. An informatics approach identifying markers of chemosensitivity in human cancer cell lines. Cancer Res. 2000;60(21):6101–6110. http://www.ncbi.nlm.nih.gov/pubmed/11085534. Accessed June 10, 2019.

67. Albeck JG, Burke JM, Aldridge BB, Zhang M, Lauffenburger DA, Sorger PK. Quantitative Analysis of Pathways Controlling Extrinsic Apoptosis in Single Cells. Mol Cell. 2008;30(1):11–25. doi:10.1016/j.molcel.2008.02.012

68. Hong J-Y, Kim G-H, Kim J-W, et al. Computational modeling of apoptotic signaling pathways induced by cisplatin. BMC Syst Biol. 2012;6(1):122. doi:10.1186/1752-0509-6-122

69. Bagci EZ, Sen SM, Camurdan MC. Analysis of a mathematical model of apoptosis: individual differences and malfunction in programmed cell death. J Clin Monit Comput. 2013;27(4):465–479. doi:10.1007/s10877-013-9468-z

70. Rehm M, Huber HJ, Dussmann H, Prehn JHM. Systems analysis of effector caspase activation and its control by X-linked inhibitor of apoptosis protein. EMBO J. 2006;25(18):4338–4349. doi:10.1038/sj.emboj.7601295

71. Hui KK-W, Dojo Soeandy C, Chang S, et al. Cell-based high-throughput screen for small molecule inhibitors of Bax translocation. J Cell Mol Med. 2019;23(3):1784–1797. doi:10.1111/jcmm.14076

72. Nakka VP, Gusain A, Mehta SL, Raghubir R. Molecular mechanisms of apoptosis in cerebral ischemia: multiple neuroprotective opportunities. Mol Neurobiol. 2008;37(1):7–38. doi:10.1007/s12035-007-8013-9

73. Erekat NS. Apoptosis and its Role in Parkinson’s Disease. In: Parkinson’s Disease: Pathogenesis and Clinical Aspects. Codon Publications; 2018:65-82. doi:10.15586/codonpublications.parkinsonsdisease. 2018.ch4

74. Tatton NA. Increased Caspase 3 and Bax Immunoreactivity Accompany Nuclear GAPDH Translocation and Neuronal Apoptosis in Parkinson’s Disease. Exp Neurol. 2000;166(1):29–43. doi:10.1006/exnr.2000.7489

75. Ma C, Pan Y, Yang Z, et al. Pre-administration of BAX-inhibiting peptides decrease the loss of the nigral dopaminergic neurons in rats. Life Sci. 2016;144:113–120. doi:10.1016/j.lfs.2015.11.019

76. Chen M, Inestrosa NC, Ross GS, Fernandez HL. Platelets Are the Primary Source of Amyloid β-Peptide in Human Blood. Biochem Biophys Res Commun. 1995;213(1):96–103. doi:10.1006/BBRC.1995.2103

77. Eberhardt O, Coelln R V, Kugler S, et al. Protection by synergistic effects of adenovirus-mediated X-chromosome-linked inhibitor of apoptosis and glial cell line-derived neurotrophic factor gene transfer in the 1-methyl-4-phenyl-1,2,3,6-tetrahydropyridine model of Parkinson’s disease. J Neurosci. 2000;20(24):9126–9134. doi:10.1523/JNEUROSCI.20-24-09126.2000

78. Palacino JJ, Sagi D, Goldberg MS, et al. Mitochondrial dysfunction and oxidative damage in parkin-deficient mice. J Biol Chem. 2004;279(18):18614–18622. doi:10.1074/jbc.M401135200

79. Ben-Shachar D, Zuk R, Glinka Y. Dopamine Neurotoxicity: Inhibition of Mitochondrial Respiration. J Neurochem. 2002;64(2):718–723. doi:10.1046/j.1471-4159.1995.64020718.x

80. Klein C, Westenberger A. Genetics of Parkinson’s Disease. Cold Spring Harb Perspect Med. 2012;2(1):a008888. doi:10.1101/CSHPERSPECT.A008888

81. Singh S, Kumar S, Dikshit M. Involvement of the mitochondrial apoptotic pathway and nitric oxide synthase in dopaminergic neuronal death induced by 6-hydroxydopamine and lipopolysaccharide. Redox Rep. 2010;15(3):115–122. doi:10.1179/174329210X12650506623447

82. Lucantoni F, Lindner AU, O’Donovan N, Düssmann H, Prehn JHM. Systems modeling accurately predicts responses to genotoxic agents and their synergism with BCL-2 inhibitors in triple negative breast cancer cells. Cell Death Dis. 2018;9(2):42. doi:10.1038/s41419-017-0039-y

83. Lindner AU, Salvucci M, Morgan C, et al. BCL-2 system analysis identifies high-risk colorectal cancer patients. Gut. 2017;66(12):2141–2148. doi:10.1136/gutjnl-2016-312287

84. Fey D, Aksamitiene E, Kiyatkin A, Kholodenko BN. Modeling of Receptor Tyrosine Kinase Signaling: Computational and Experimental Protocols. In: Methods in Molecular Biology (Clifton, N.J.). Vol 1636.; 2017:417–453. doi:10.1007/978-1-4939-7154-1_27

85. Röst HL, Rosenberger G, Navarro P, et al. OpenSWATH enables automated, targeted analysis of data-independent acquisition MS data. Nat Biotechnol. 2014;32(3):219–223. doi:10.1038/nbt.2841

86. Rajapakse VN, Luna A, Yamade M, et al. CellMinerCDB for Integrative Cross-Database Genomics and Pharmacogenomics Analyses of Cancer Cell Lines. iScience. 2018;10:247–264. doi:10.1016/j.isci.2018.11.029

