## Supplementary figures and images for "BAX and SMAC Regulate Bistable Properties of the Apoptotic Caspase System"

### Supplemental Data 2

#### Supplement figure 1

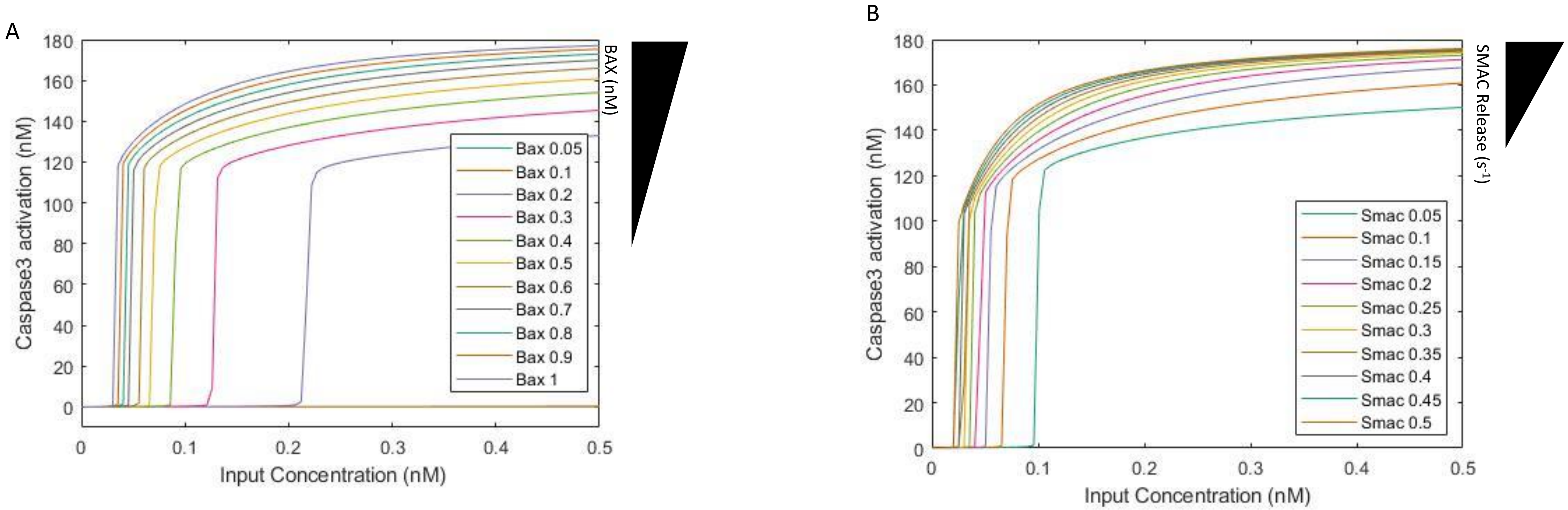

### Supplemental Data 3

**Supplement Figure 2**

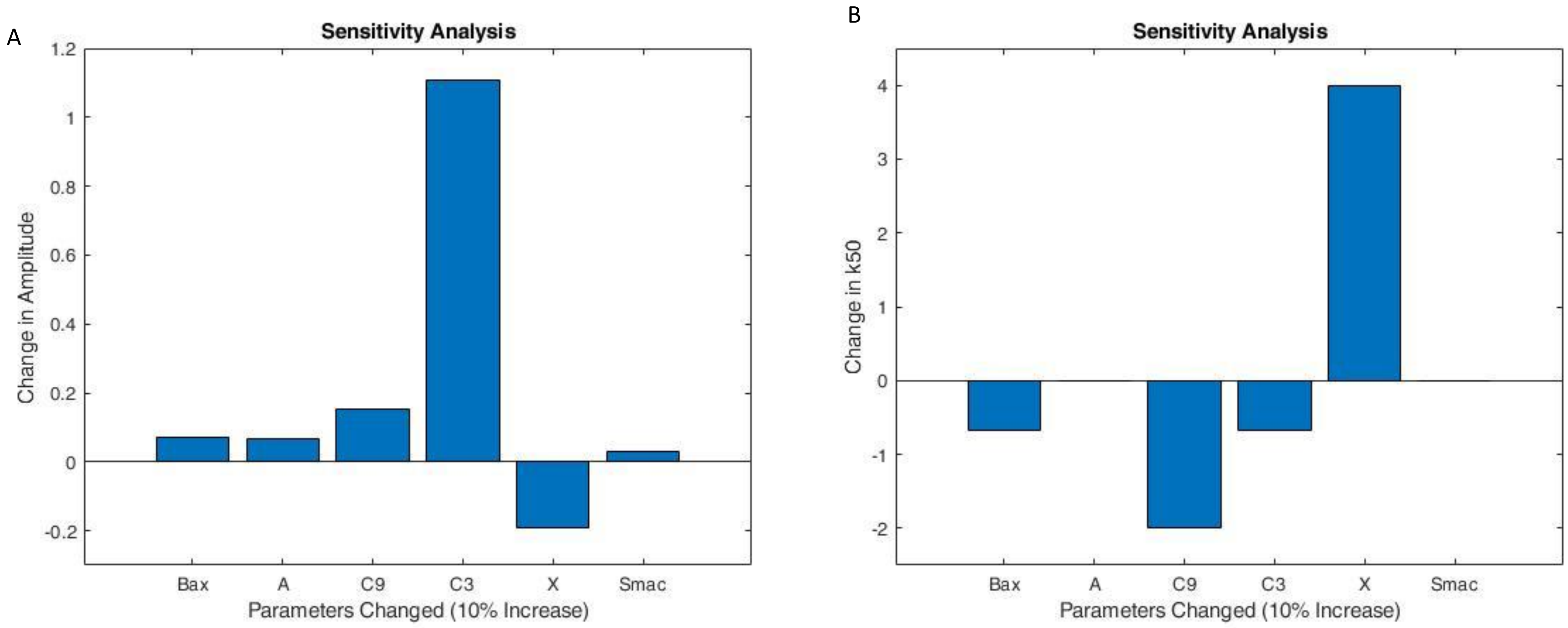

### Supplementary Figure 1

**Supplemental Figure 3**

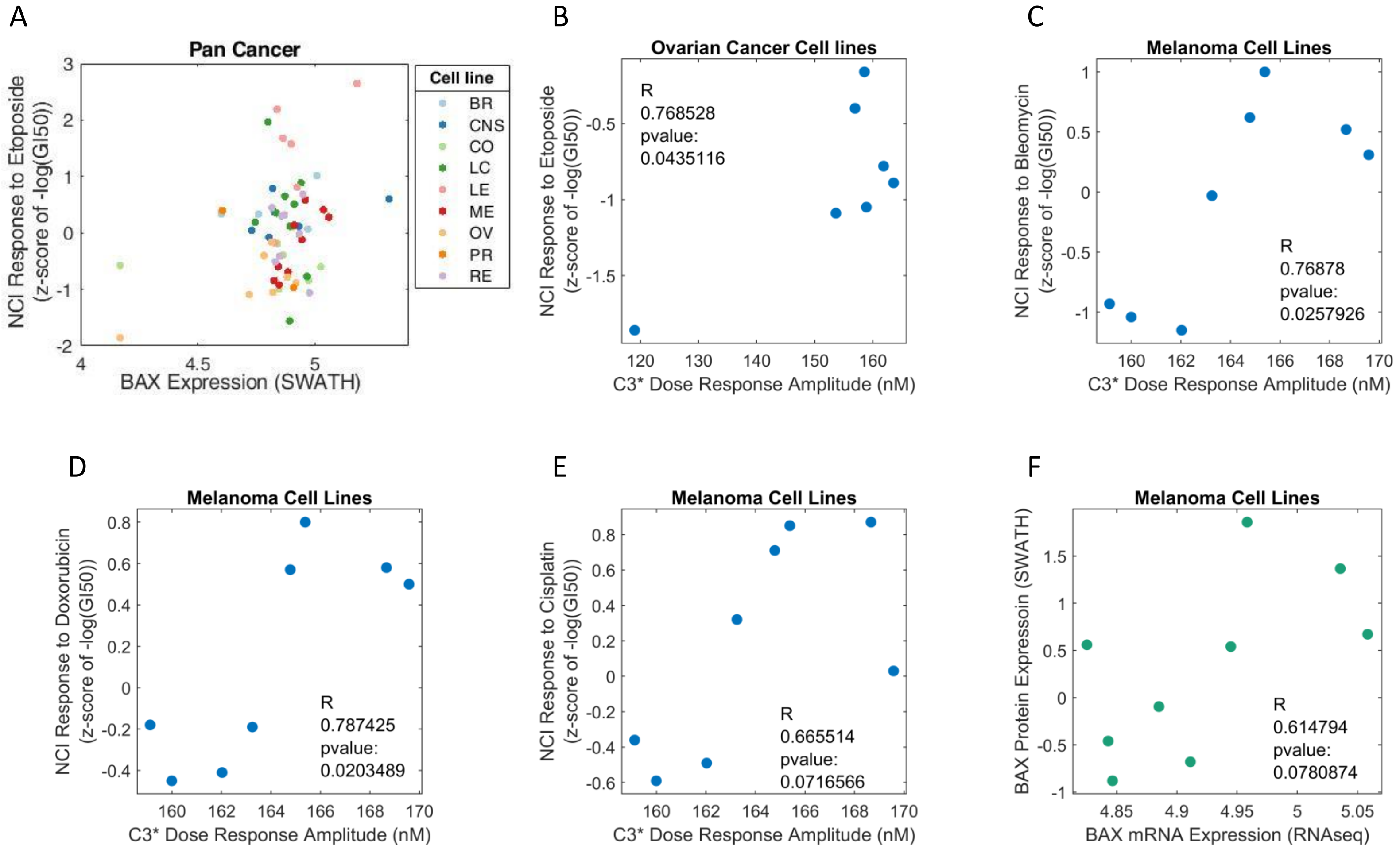
