## Supplemental Data 1 for "BAX and SMAC Regulate Bistable Properties of the Apoptotic Caspase System"

### Supplementary Information

Supplementary Figure 1.

(A) Dose response simulations of caspase 3 activation for a range of BAX initial concentrations (BAX0) (0.1-1nM). Nominal BAX0 is 0.5nM. The concentration of input stimulus required to activate caspase 3 decreased with increase in BAX0. SMAC remained as in the nominal model for all simulations with SMAC release rate =0.1s^-1^.
(B) Dose response simulations for a range of SMAC release rates (0.05-0.5s-1). Nominal SMAC release rate is 0.1 s^-1^. Upon increase in SMAC release rate, a decrease in concentration of input stimulus required to activate caspase 3 is observed.

Supplementary Figure 2. Sensitivity Analysis of Model Parameters.

A sensitivity analysis was carried out for the model species. Parameters were increased by 10%, model simulations were run and results were normalised by the change in parameter. (A) It emerged that a 10% increase in Caspase 3 concentration had the highest impact on C3 amplitude. (B) A 10% increase in XIAP concentration had the highest impact on C3 activation threshold.

Supplementary Figure 3. Cell Line Specific Simulations and Correlation with Drug response.

A. Pan cancer BAX levels (protein SWATH) positively correlate with response to Etoposide (C=0.307, pval=0.019). B. Ovarian cancer cell line specific simulation C3 activation amplitude positively correlates with ovarian cell line response to Etoposide (C=0.77, pval=0.04). C-E. Melanoma cell line specific simulation C3 amplitude demonstrates positive correlations with alternative DNA damaging agents: Bleomycin, Doxorubicin, Cisplatin. F. BAX mRNA expression and protein (SWATH) levels are positively correlated (C=0.615, pval=0.078).
